# tICA-Metadynamics for Identifying Slow Dynamics in Membrane Permeation

**DOI:** 10.1101/2023.08.16.553477

**Authors:** Myongin Oh, Gabriel C. A. da Hora, Jessica M. J. Swanson

## Abstract

Molecular simulations are commonly used to understand the mechanism of membrane permeation of small molecules, particularly for biomedical and pharmaceutical applications. However, despite significant advances in computing power and algorithms, calculating an accurate permeation free energy profile remains elusive for many drug molecules because it can require identifying the rate-limiting degrees of freedom (i.e., appropriate reaction coordinates). To resolve this issue, researchers have developed machine learning approaches to identify slow system dynamics. In this work, we apply time-lagged independent component analysis (tICA), an unsupervised dimensionality reduction algorithm, to molecular dynamics simulations with well-tempered metadynamics to find the slowest collective degrees of freedom of the permeation process of trimethoprim through a multicomponent membrane. We show that tICA-metadynamics yields translational and orientational collective variables (CVs) that increase convergence efficiency ∼1.5 times. However, crossing the periodic boundary is shown to introduce artefacts in the translational CV that can be corrected by taking absolute values of molecular features. Additionally, we find that the convergence of the tICA CVs is reached with approximately five membrane crossings, and that data reweighting is required to avoid deviations in the translational CV.

## INTRODUCTION

Lipid membranes function as the physical barrier that controls the exchange of matter, energy, and information between cells and organelles in biological systems. Membrane permeation of small molecules is often an activated process that takes place on timescales inaccessible by standard molecular simulations. The dynamics of large biomolecular systems is governed by a complex high-dimensional free energy landscape characterized by a hierarchy of energy barriers.^1-3^ Transitions between metastable states become rare when they are separated by large energy barriers while thermal fluctuations are the only driving force for barrier crossing.^4^ These kinetic bottlenecks restrict the timescale that can be explored (which is generally in the range of microseconds or shorter) by conventional molecular dynamics (MD) simulations, thereby introducing substantial statistical errors in the measurement of structural, thermodynamic, and kinetic properties. To circumvent this sampling issue, many of the existing computational methods used to evaluate membrane permeation rely on enhanced free energy sampling techniques.^5-9^ Impressively, membrane permeation has been directly observed for a few small molecules in canonical MD simulations on the nano- and microsecond timescales. For instance, Krämer *et al*.^10^ performed unbiased MD simulations to evaluate the permeability coefficients of oxygen, water, and ethanol using counting methods and maximum likelihood estimation for the inhomogeneous solubility-diffusion (ISD) model.^6, 11-12^ They found that counting methods^13^ yield nearly model-free estimates for all the three permeants, whereas the ISD model causes large uncertainties for water due to insufficient sampling and overestimates for ethanol due to collective effects in the membrane.^10, 14^ For larger molecules with slower permeation, however, enhanced sampling can be essential to increase the occurrence of the slowest dynamical motions that enable permeation. This can be done by biasing the potential energy surface or altering the probability density of sampled conformations.^15-16^ For example, an external bias potential can be added to the Hamiltonian (as in umbrella sampling and metadynamics (MetaD)), or the system can be coupled to higher temperatures (as in replica exchange MD) to effectively reduce energy barriers and thus sample transition regions, or the transition ensemble can be selectively sampled with path sampling approaches, such as transition interface sampling or its combination with replica-exchange with or without memory effects.^14, 17^

The success of enhanced free energy sampling methods involving a Hamiltonian bias requires the selection of a proper set of collective variables (CVs). CVs are user-defined functions of atomic (Cartesian) coordinates that provide a low-dimensional projection of conformational phase space while, in principle, retaining ‘important’ information. Dimensionality reduction is an essential consequence of our inability to work in, or visualize, high-dimensional spaces. Only in a reduced number of dimensions (typically 2 to 4) can we define effective bias potentials to alter dynamics, visualize complex free energy landscapes, efficiently sample conformational phase space that crosses high energy barriers, and simplify simulation data for noise omission and better inference (*i*.*e*., escape the curse of dimensionality).^18^ Ideally, CVs are translationally and rotationally invariant, include all relevant slow molecular motions, and distinguish local minima (metastable states) and activation barriers (transition states).^19-22^ If properly obtained, CVs provide lower variance estimators of the properties of interest when the system diffuses on the free energy surface (or potential of mean force, PMF) spanned by these CVs. However, it is highly nontrivial to intuit good CVs for complex systems (large biomolecules in particular), and accelerating dynamically irrelevant CVs may give rise to inaccurate profiles and decreased efficiency compared to unbiased MD.^21^ In recent years, the availability of large datasets (obtained directly from unbiased and biased simulations) in conjunction with advances in computing hardware and algorithms has led to the automated design of CVs inspired by machine learning, data science, and information theory. A comprehensive review of machine learning approaches for CV discovery is presented in refs. [^4,18,21, 23-24^].

Data-driven CVs are typically coincident with collective degrees of freedom that either have high variance or evolve slowly. High-variance CVs can be identified by the principal component analysis (PCA), an unsupervised linear transformation that finds a subspace that maximally preserves the configurational variance contained within a molecular simulation trajectory.^21^ Since the orthogonal eigenvectors, or the principal components (PCs), represent large-amplitude collective motions in terms of variance, they are often called essential dynamics^25-30^ and used as collective variables for enhanced sampling. However, a major problem inherent in PCA is that it is not generally guaranteed that large-amplitude motions are associated with the slow motions that enable transitions between metastable states. For example, Naritomi and Fuchigami^31^ found that a closure motion of the lysine-arginine-ornithine binding protein described by the largest-amplitude mode determined by PCA does not represent the slowest mode, a twist motion that takes place on a timescale of tens of nanoseconds. Generally, identifying slow motions, rather than large-amplitude motions, is essential to address the sampling problem. tICA, initially introduced as a signal decomposition algorithm^32^, is an unsupervised linear transformation that finds a subspace that maximally preserves the kinetic content (*i*.*e*., minimizes the loss of kinetic information) by maximizing the autocorrelation function.^21, 33-35^ The resulting eigenvectors, or independent components (ICs), represent the slowest-relaxing degrees of freedom in time-series data and approximate the eigenfunctions of the underlying Markovian dynamics. In other words, tICA provides the optimal linear approximation to the variational approach to conformational dynamics, a systematic approach for modeling the slow parts of Markov processes by approximating the dominant eigenfunctions of the MD propagator (or transfer operator).^36-39^ Concisely, tICA is a special case of the linear variational approach which uses mean-free input descriptors as a basis set and empirical estimates of their covariance matrices in an eigenvalue problem.^33, 36^ More details on tICA can be found in Methods.

Once slowly decorrelating modes are discovered, they can be biased in CV-based enhanced sampling techniques to accelerate the occurrence of rare events. In tICA-MetaD, tICA is performed on MD simulation trajectories to identify the ICs which are then directly used as CVs in MetaD to obtain highly diffusive behavior in CV space and fast convergence of PMF calculations. For instance, Sultan and Pande^40^ applied linear and nonlinear tICA-MetaD on unbiased MD simulations of alanine dipeptide and bovine pancreatic trypsin inhibitor to explicitly sample their slowest modes. In principle, the combined method can drive slow transitions even when no such transitions take place in the original unbiased simulations. However, this only happens when the slow motions captured in the original simulations are the same as those that enable the sought-after transition(s). This can be a major limitation of their method since it requires unbiased sampling of the relevant slow transitions (*e*.*g*., via very long aggregate MD sampling or enough MD runs starting from high and low free energy states), which is often challenging or even unfeasible depending on the system.^40^ A clear example of this would be the permeation of a highly polar molecule for which unbiased sampling never entered the hydrophobic membrane midplane.

To resolve this problem, McCarty and Parrinello^41^ proposed to start with a biased simulation with suboptimal CVs, reweight the trajectory to recover unbiased distributions (*i*.*e*., Boltzmann statistics) of the system, and perform tICA to derive slow CVs for MetaD. This CV optimization method has been applied to different systems ranging from conformational transitions of alanine dipeptide, alanine tetrapeptide^41^, and L99A T4 lysozyme^42^ to homogeneous crystallization of sodium and aluminum^43^. However, their method also faces several challenges. First, data reweighting is not a trivial task; for example, their reweighting scheme may not work properly on the trajectory generated by transition-tempered metadynamics (TTMetaD).^8, 44^ Second, reweighting does not necessarily converge accurately and rapidly to the underlying free energy depending on the reweighting technique, the enhanced sampling technique, and the stage of the simulation.^45^

Previously, we studied the molecular mechanism of membrane permeation of trimethoprim, an antibacterial agent primarily used in the treatment of urinary tract infections,^46^ using an alternative implementation of tICA-MetaD.^47^ First, we performed a short TTMetaD simulation with non-optimal CVs to generate an initial trajectory that involved at least one transition of interest (permeation). Second, we identified the slowest decaying modes using tICA without data reweighting. Lastly, we carried out a new TTMetaD simulation using a permeation CV consistent with the original bias, and the slow CV obtained from tICA. We then examined the accuracy and convergence of PMF calculations. Interestingly, the use of the tICA CVs was shown to accelerate the convergence of PMF calculations while also revealing a subtle effect of cholesterol on the permeation of the small drug molecule through the heterogeneous model membrane. Perhaps more importantly, this work brought to light several outstanding questions regarding the optimal use of tICA-MetaD. For example, it remains unclear what effect trajectory reweighting has on tICA CVs, whether or not this approach is applicable to other forms of tempering, and how many rare events are needed in the original trajectories in order to obtain consistent (converged) tICA CVs. Moreover, the influence of the periodic boundary conditions (PBCs) on tICA-like analyses of permeation has, to the best of our knowledge, been largely overlooked.

In this work, we apply several variations of tICA-MetaD to membrane permeation to address these outstanding questions. We opt for a pair of molecular features, specifically the z-positions of trimethoprim relative to the lipid bilayer, and design a straightforward yet suboptimal initial CV. This approach aims to initiate rare-event crossings to obtain trajectory data that can be fed into tICA-MetaD protocols in order to identify optimal CVs (Figure S16). The use of a simple and largely transferable initial CV (such as z-position relative to the membrane) is particularly relevant when the selection of good CVs is not known a priori. We find that tICA-MetaD identifies the translational and orientational modes as slow CVs, and the use of the machine-learned tICA CVs leads to ∼1.5 times faster convergence of our PMF calculations. However, the tICA-MetaD scheme we employed cannot be applied to TTMetaD trajectories. Also, the PBCs can be detrimental to tICA for membrane permeation trajectories. The orientational CV is not significantly affected by the PBCs, while taking absolute values of molecular features corrects the error in the translational CV. Additionally, the convergence of the tICA CVs is attained even when only five membrane crossings are included in the trajectory regardless of data reweighting, and the first eigenvector may deviate significantly from the translational CV when data reweighting is omitted. Collectively, we hope these findings will help inform future implementation best practices for tICA-MetaD applied to membrane permeation.

## METHODS

### All-atom Molecular Dynamics Simulation

We performed all-atom molecular dynamics simulations using GROMACS 2019.4^48^ patched with PLUMED 2.5.3^49^ for well-tempered metadynamics (WTMetaD) and TTMetaD. For optimal comparison, the heterogeneous bilayer system used in our previous work selected for this study, including 36 phosphatidylcholine (POPC) and 4 cholesterol molecules (*n*_POPC_: *n*_CHOL_ = 9: 1). The bilayer was placed on the *xy*-plane (*i*.*e*., with the surface normal along the *z*-axis) and solvated by 2200 water molecules in a unit cell with dimensions of approximately 3.6 nm × 3.6 nm × 9.5 nm. The PBC was then applied in every direction. CHARMM36^50^ and the CHARMM General Force Field (CGenFF)^51^ were employed to model the lipid molecules and trimethoprim, respectively. The water molecules were modeled with the TIP3P^52^ potential. We prepared the initial structure of the lipid bilayer using the CHARMM-GUI membrane builder^53^ and equilibrated it for 150 ns in water. After equilibration, we placed the drug molecule randomly into the aqueous region using PACKMOL^54^. To generate an isobaric-isothermal (NPT) ensemble of the system at 323.15 K and under 1 bar, the lipid and the water plus the drug molecule were separately coupled to two heat baths using velocity rescaling with a stochastic term^55^, while the system was coupled semi-isotropically to a Berendsen barostat^56^ such that the simulation box was rescaled every 5 ps. The cutoff distance for the short-range neighbor list was set to be 12 Å, and the neighbor list was updated every 40 steps. Fast smooth Particle-Mesh Ewald (SPME)^57^ was used to treat long-range electrostatic interactions. All covalent bonds including hydrogen atoms were constrained by linear constraint solver (LINCS)^58^, and the integration timestep was set to be 2 fs. The initial velocities were randomly sampled from a Maxwell-Boltzmann distribution at 323 K which is well above the transition temperature of the lipids. We generated the images from the simulations and analyzed the MD trajectories using Visual Molecular Dynamics (VMD) version 1.9.3^59^. Details of each simulation are described in Table 1.

**Table 1.**
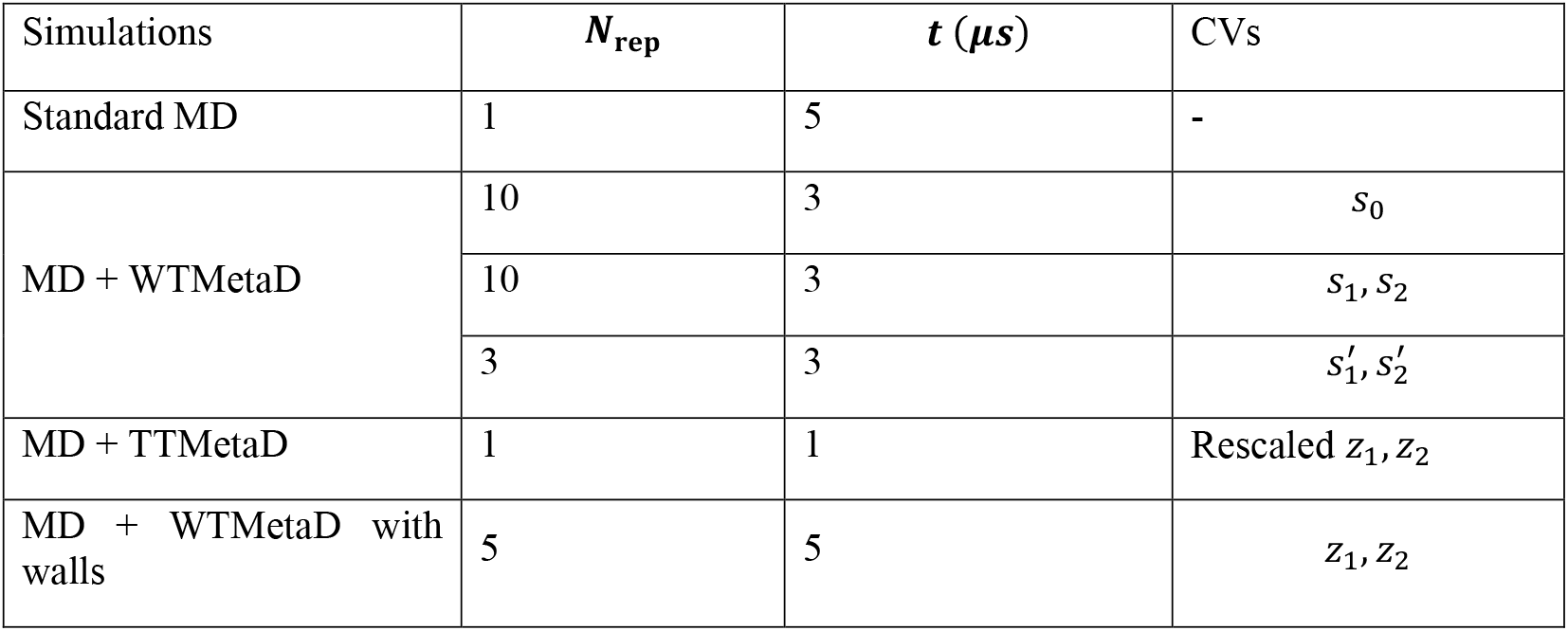
Summary of simulations showing the type of simulation, the number of replicas (***N***_**rep**_), and the length (***t***) of each replica, and the CVs biased. *z*_1_ and *z*_2_ are molecular descriptors, *s*_0_ is the suboptimal CV, *s*_1_ and *s*_2_ are the optimal CVs, and 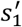 and 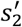 denote the CVs obtained with |*z*_1_| and |*z*_2_|.

To investigate the effect of the PBC on tICA CVs, we applied lower and upper walls that limit the phase space accessible to the system during the simulation. In PLUMED 2.5.3, these walls are implemented in the form of restraining potentials *η*(***s***) given by

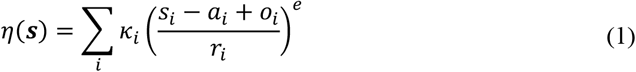

where *κ*_*i*_, *s*_*i*_, *a*_*i*_, *o*_*i*_, and *r*_*i*_ denote a force constant, a CV, the position of the wall, an offset, and a rescaling factor, respectively, and *e* is the exponent that determines the power law. In our simulations, we set *o*_*i*_ = 0, *r*_*i*_ = 1, and *e* = 2 to realize harmonic restraints and added both upper and lower walls on *z*_1_ and *z*_2_ (*i*.*e*., suboptimal CVs) at 3.6 nm and −3.6 nm with *κ*_*i*_ = 1500 kJ/mol.

### Well-tempered Metadynamics

WTMetaD^19, 60-61^ is an enhanced sampling technique that discourages the system from revisiting configurations that have previously been sampled in the CV space by periodically adding to the Hamiltonian of the system small repulsive Gaussians whose amplitude decreases exponentially as the simulation progresses. The sampling of rare events is accelerated due to ergodic and diffusive motion of the system along the selected CVs. The instantaneous bias potential *V*(*s, t*) at point *s* in CV space and at time *t* is determined by the sum of Gaussian hills deposited over the past trajectory of the system:

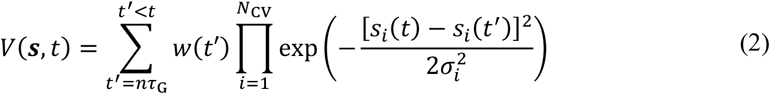

where *σ*_*i*_ is the width of the Gaussian hill for *s*_*i*_, the *i* ^th^ CV. *N*_CV_ is the number of CVs, *τ*_G_ is the stride of Gaussian deposition, and *w* is the adaptive height of the Gaussian hill given by

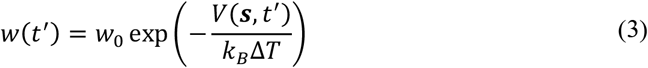

where *w*_0_ is the initial height of the Gaussian hill, *k*_*B*_ is the Boltzmann constant, and Δ*T* is a parameter that tunes how quickly the Gaussian height is reduced. Ideally, Δ*T* should be proportional to the free energy barrier to be crossed, which is generally not known in advance. If the parameter is too small, then the system fails to escape local minima; if the parameter is too large, unphysical instability can be introduced causing significant errors in the PMF. The adjustment of the Gaussian deposition allows smooth convergence and tunable error of the bias potential so that the unbiased free energy *F*(*s*) of the system can be estimated as the limit

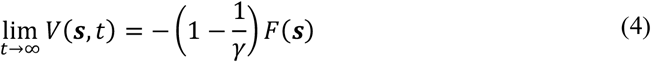

where γ is a bias factor at temperature *T* given by

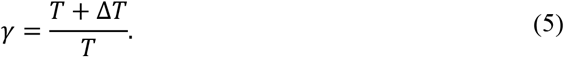

An advantage of WTMetaD is that after a transient time, the simulation enters a quasi-stationary limit in which the expectation value of any observable ⟨*O*(***R***)⟩ can be estimated as a running average according to

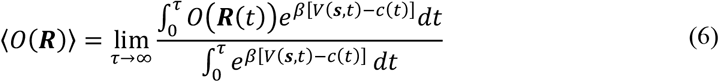

where ***R***(*t*) is atomic position at time *t, β* is the reciprocal of the product of *k*_*B*_ and *T*, and *c*(*t*) is a time-dependent bias offset defined as

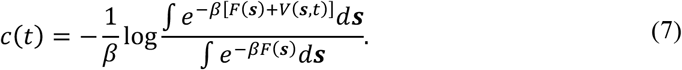

The function *c*(*t*) asymptotically tends to the reversible work done on the system by the external bias.

In our WTMetaD simulations, the Gaussian bias was deposited every 500 steps with a height of 0.05 kJ/mol and a width of 0.2 nm. The height of the Gaussian hill was tempered with a bias factor of 15. Multiple replicas were prepared, initialized randomly, and run for 3-5 *μ*s, and the resulting PMFs were obtained by averaging over replicas. To generate 1D PMFs, we further symmetrized the free energy profiles with respect to the center of the horizontal axis. The minimum free energy paths (MFEPs) on 2D PMFs were calculated with a zero-temperature string method^62-63^ which represent the most probable transition paths in the ensemble of the permeation processes. The 1D PMF was directly obtained by using the average MFEP from the independent replicas. Error bars were computed using the standard errors across replicas.

### Time-lagged Independent Component Analysis

The microstate of a molecular system can be defined by a few degrees of freedom, or features, which are selected prior to tICA and used as input to tICA. Our selection of features was motivated by previous work in which the *z*-position of the center of mass (COM) of each ring of trimethoprim relative to that of the lipid membrane was shown to reduce the total simulation time required for convergence of the permeation PMF.^64-65^ The positions were denoted by *z*_1_ for the trimethoxybenzyl (TMB) group and *z*_2_ for the diaminopyrimidine (DAP) group (Fig. 1(c)). Also, the two CVs were shown to capture orientational changes of the drug molecule relative to the membrane surface normal. However, it is not guaranteed that the two CVs are (1) associated with slow transitions or (2) optimal compared to many other degrees of freedom that mediate the permeation process. In addition, we chose these simple suboptimal CVs as features for better interpretability of optimal CVs and comparison with previous results.

**Figure 1.**
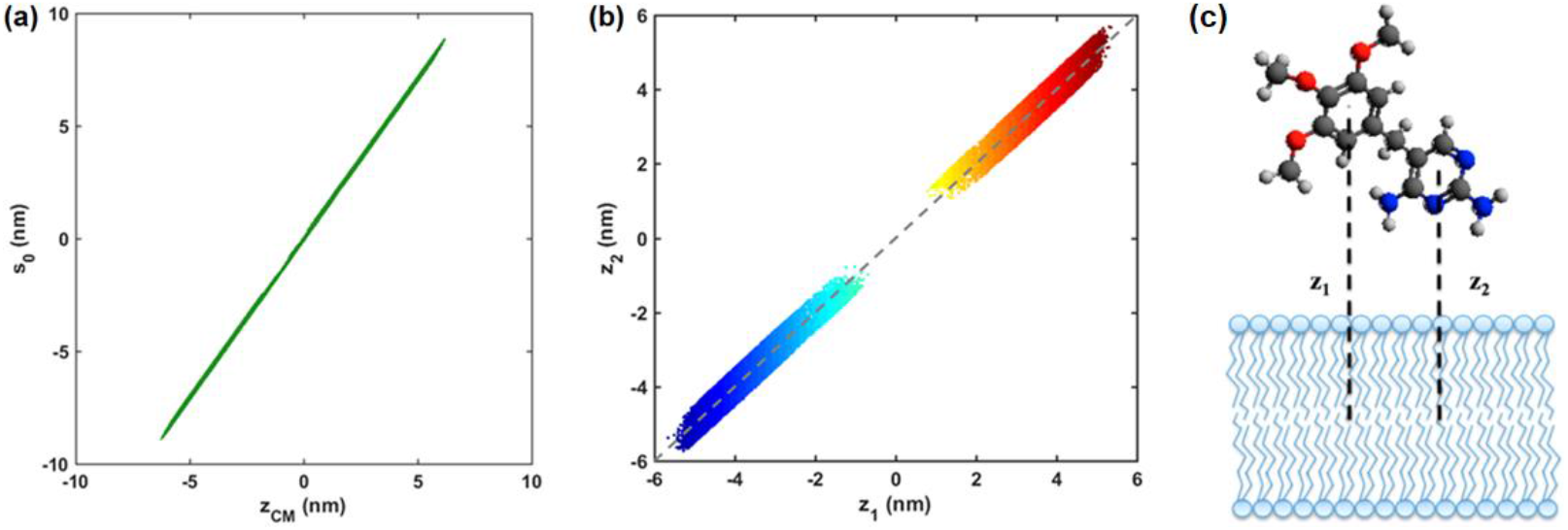
(a) Linear relationship between the suboptimal CV *s*_0_ and the *z*-position *z*_*CM*_ of the COM of trimethoprim with respect to that of the lipid bilayer. (b) Scatter plot displaying all the (*z*_1_, *z*_2_) pairs accessed in 5μs unbiased MD simulation. The red and yellow colors indicate the region above the bilayer, and the dark and light blue colors represent the region below the bilayer. The gap between the two regions signifies the interior of the membrane. The dashed gray line shows the first PC, *s*_0_. (c) Molecular structure of trimethoprim and definitions of the molecular features *z*_1_ and *z*_2_.

Once the covariance matrices central to tICA were computed, these features were then linearly combined to generate two CVs which optimally describe the slow modes of the system by solving a generalized eigenvalue problem. Robustness of tICA CVs was examined over a range of lag times from 1 to 9 ns. Finally, tICA CVs were biased as new CVs in WTMetaD simulations to investigate any significant improvement in sampling efficiency. To understand the effect of biasing on tICA CVs, we applied the method to both biased and unbiased subtrajectories which involve different numbers of drug permeation events. We obtained unbiased trajectories from biased ones through a reweighting scheme as described in the next section.

Here, we provide a brief introduction to the method as the theoretical background of tICA can be found in many other works. tICA is an unsupervised dimensionality reduction algorithm designed to find a maximally slow subspace ***U*** via a linear transformation of molecular features ***χ*** by maximizing the autocorrelation function of their projections. In other words, it finds the degrees of freedom where correlation decays slowly and provides an optimal linear solution to the variational approach to conformational dynamics that approximates the slow components of reversible Markov processes. Mathematically, tICA determines the slowest independent collective degrees of freedom ***u***_*i*_ onto which the projections *s*_*i*_(*t*) = ***u***_*i*_ *·* ***χ***(*t*) have the largest autocorrelation function

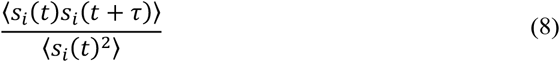

for a chosen lag time *τ*. Equivalently, tICA maximizes the ratio *J*(***U***)

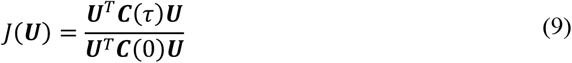

with respect to ***U***= [***u***_1_, ⋯, ***u***_*d*_]. This is equivalent to finding the solution of the generalized eigenvalue problem

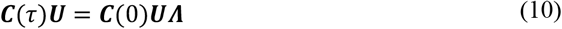

where *τ* is the lag time, ***C*** is the covariance matrix (*i*.*e*., ***C***(τ) = ⟨***χ***(*t*)***χ***^*T*^(*t* + *τ*)***⟩***_*t*_ where the data ***χ*** is mean-free), ***U*** is the eigenvector matrix that contains linearly independent components (ICs), and ***Λ*** is a diagonal eigenvalue matrix that contains autocorrelations. The dominant ICs are used to reduce the dimensionality of the phase space and capture the slowest modes of the molecular system. A direct estimation of ***C***(τ) from finite trajectories generally yields an asymmetric matrix and complex-valued eigenvectors and eigenvalues which are inconsistent with reversible molecular dynamics. To circumvent this problem, a symmetric estimator is often used to approximate the covariance matrices by averaging over all pairs (***x***_*t*_, ***x***_*t*+τ_) and the reverse (***x***_*t*+τ_, ***x***_*t*_), which is equivalent to averaging time-forward and time-reversed trajectories, where ***x*** is a set of mean-free molecular coordinates^34^.

tICA suffers from one significant pitfall: the choice of a proper value of *τ* which determines which kinetic processes are considered to construct the new subspace. Ideally, *τ* must be sufficiently long that the dynamics described by ICs satisfies the Markovian property of being memoryless, but also sufficiently short that the significant dynamics are not neglected. Currently, there is no recipe to choose an appropriate value of the parameter^18^. Therefore, ICs are ideally robust within a range of *τ* smaller than their timescales, but it is practically inevitable that they change to some extent as the parameter value changes^66^. In this work, we examine the components of the eigenvectors as a function of *τ*, as done previously by Parrinello and colleagues^41, 67^.

### Reweighting Algorithm

Information about the unbiased state of the system can be recovered directly from a metadynamics simulation by means of a reweighting procedure^45^. We approximated the unbiased slow modes of the system from our WTMetaD trajectories using the reweighting scheme proposed by McCarty and Parrinello.^41^ Here, we summarize the key steps involved in their reweighting procedure. For the column vector ***χ***(*t*) of the molecular features at time *t*, the two covariance matrices in Eq. 10 are calculated as follows:

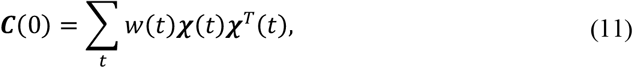

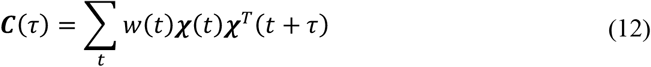

where the statistical weight *w*(*t*) of the configuration at time *t* is given by

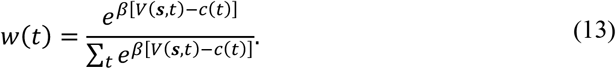

In the case where walls are employed, the potentials are not included in the calculation of the statistical weights. Furthermore, the time scale in biased simulations needs to be properly rescaled as follows:

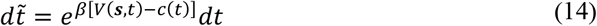

so that *τ* is given by the sum of the rescaled time steps

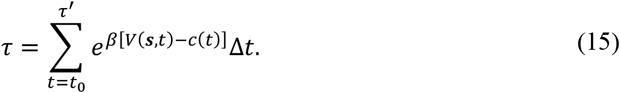

If *N* molecular features are chosen to define the basis functions, then *M* eigenvectors that correspond to the slowest relaxation times (*i*.*e*., largest eigenvalues) are selected to be biased as optimal CVs with WTMetaD:

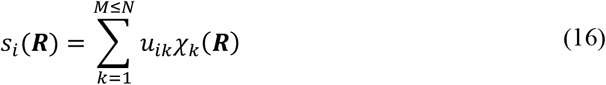

where ***u***_*i*_ is the *i*^th^ eigenvector in ***U***(or the set of expansion coefficients) in Eq. 10.

### Contact Analysis

Contact analysis was performed for the DAP and TMB moieties interacting with different components of the system, including water, cholesterol, POPC, and the choline-phosphate (PO4) group or ester region of POPC (Figure S11). A contact was defined as any atom present within a cutoff radius of 3.5 Å of the reference group. The analysis was performed using the MDAnalysis Python package^68^. The same groups were used to calculate the average interaction energy using the *gmx energy*.

## RESULTS AND DISCUSSION

### Application of tICA-MetaD to Membrane Permeation Simulations

We employed the tICA-MetaD scheme (Figure S16) proposed by McCarty and Parrinello^41^ to identify unbiased slow CVs for the passive diffusion of trimethoprim across a model heterogeneous lipid membrane composed of POPC and cholesterol. The procedure starts from the design of any arbitrary, suboptimal CV *s*_0_ that enhances the occurrence of rare events and distinguishes different metastable and transition states. Here, we defined *s*_0_ as a sum of *z*_1_ and *z*_2_ (*i*.*e*., the molecular descriptors, Fig 1c) with equal weights, namely, *s*_0_ = 0.7071*z*_1_ + 0.7071*z*_2_. Mathematically, this is simply the unit vector of (1,1) in the descriptor space of *z*_1_ and *z*_2_. Physically, *s*_0_ carries the same information as *z*_CM_ (*i*.*e*., the *z*-position of the COM of the drug molecule relative to that of the lipid bilayer) but is rescaled; when we plot *s*_0_ as a function of *z*_CM_ using one of the WTMetaD trajectories biasing *s*_0_ (see the following paragraph), we observe a linear relationship between the two variables, namely, *s*_0_ = 1.4153*z*_CM_ with R^2^ = 0.9997 (Fig. 1(a)). Interestingly, *s*_0_ is also the first PC that retains 99.84 % of the total variance when PCA is applied to the (*z*_1_, *z*_2_) points obtained from a 5 *μ*s-long unbiased trajectory of the system (Fig. 1(b)).

Next, we performed WTMetaD simulations that bias *s*_0_ and thus induce multiple permeation events. In this step, we generated 10 different biased trajectories for reliable statistical analysis, and each trajectory was 3-*μ*s long. Fig. 2(a) shows how the suboptimal CV *s*_0_ evolves with time in one of the biased trajectories. The dashed lines indicate the average positions of the phosphorous atoms of POPC. It is clear that *s*_0_ induces transitions between the local minima. The average rate of complete permeation (*i*.*e*., mean recrossing frequency) was calculated to be 3.3 ± 1.3 *μs*^−1^. (To determine the average rate, we divided the total number of membrane crossings by the total length of the simulation.) Figure 2(d) shows the difference γ(*t*) between the instantaneous bias potential *V*(*s, t*) (Fig. 2(b)) and the bias offset *c*(*t*) (Fig. 2(c)) as a function of time, which determines the statistical weight of each time frame in the reweighting procedure. The gray region in Fig. 2(c) indicates the transient time to be eliminated for tICA.

**Figure 2.**
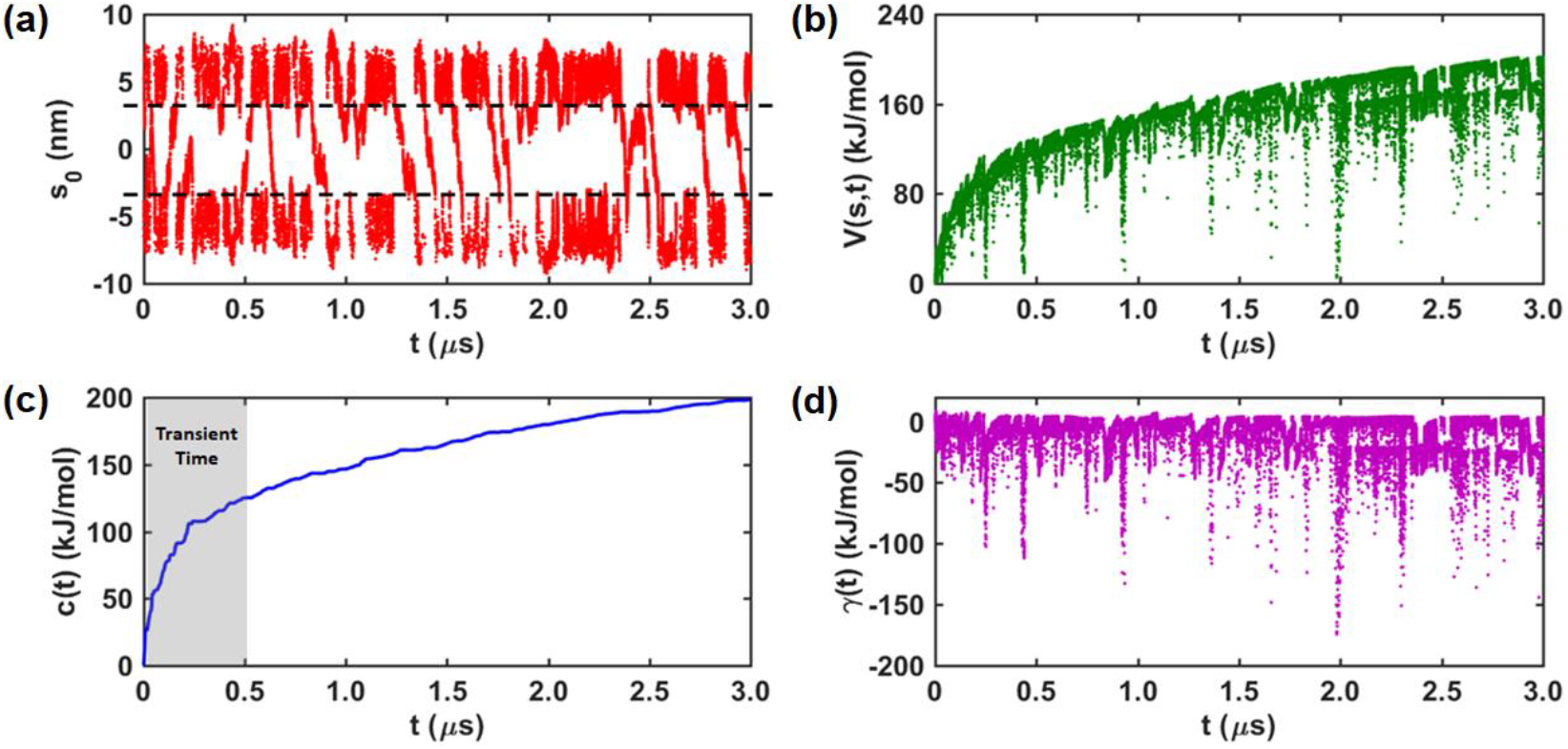
Time evolution of (1) the suboptimal CV *s*_0_, (2) the instantaneous potential *V*(*s, t*), (3) the bias offset *c*(*t*), and (d) the difference γ(*t*) between *V*(*s, t*) and *c*(*t*) in one of the WTMetaD simulations we performed. The dashed black lines in (a) indicate the average position of the phosphorous atoms of the lipid bilayer. The gray region in (c) indicates the transient time to be removed for tICA.

The average 1D PMF for the permeation process is presented in Fig. 3. Two local minima are found at *s*_0_ = ±2.3 nm, which are separated from the aqueous regions (corresponding to the regions where *s*_0_ is in the range of [−5, −4] nm or [4, 5] nm) by small energy barriers located at *s*_0_ = ±3.3 nm. Relative to the bulk region, the depth of the minima is ∼1.0 kJ/mol, and the height of the barriers is approximately 4.0 kJ/mol. The system reaches the local minima when the TMB group resides in the hydrophobic region of lipid tails while the DAP group interacts with the polar headgroups (Fig. 3(a and c) and Fig. 5(c)). A large, broad energy barrier of 32.4 kJ/mol, which connects the two local minima, takes place at *s*_0_ = 0 nm. The energy barrier arises when trimethoprim moves and flips inside the hydrophobic core of the lipid bilayer. *s*_0_ can describe the translational motion but not the orientational change of the drug molecule. For example, we observed from unbiased simulations less populated conformations in which the DAP group dwells in the hydrophobic region while the TMB group is exposed to the polar region of the lipid bilayer, but the corresponding values of *s*_0_ to those conformations are not distinguishable. From this observation, we anticipate that the optimal slow CVs should account for both translational and orientational modes of the permeant.

**Figure 3.**
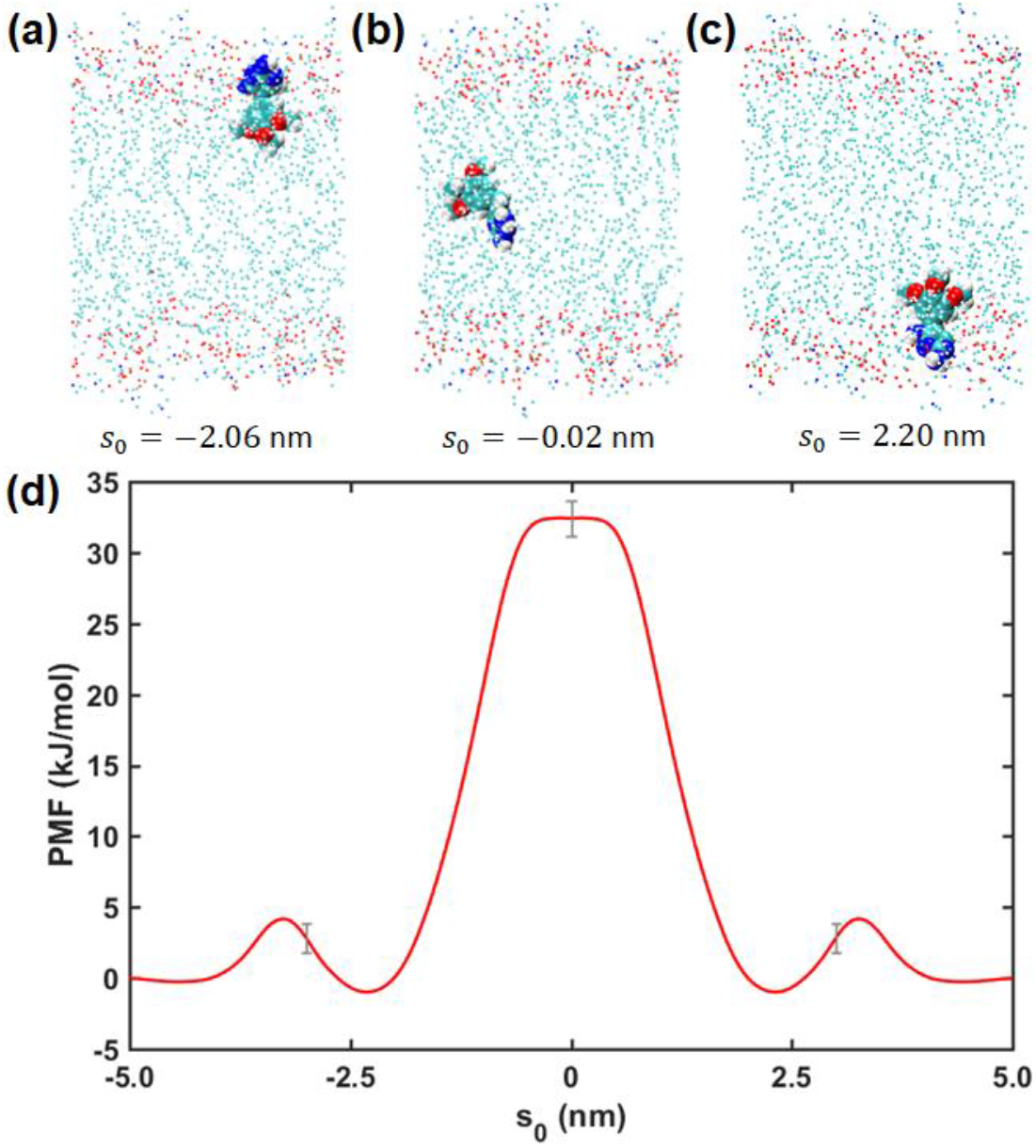
(a-c) Typical snapshots of the system in the local minima and at the transition state with the corresponding values of *s*_0_. (d) Average 1D PMF along the suboptimal CV *s*_0_ for the permeation process of trimethoprim through the lipid bilayer. The free energy in bulk water was set to zero, and the error bars indicate the standard deviations for arbitrarily selected points.

To identify the unbiased slow modes, we reweighted the biased trajectories and performed tICA on them as proposed by McCarty and Parrinello.^41^ The results from tICA on the first reweighted trajectory are summarized in Fig. 4. The red and yellow curves indicate the first and second components 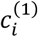 of the first eigenvector (or IC) in Fig. 4(a); likewise, the green and chartreuse curves represent the first and second components 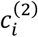 of the second eigenvector in Fig. 4(b). The components of the eigenvectors are fairly stable with respect to *τ*. The slow CVs are expressed as a linear combination of the chosen molecular features: *e*.*g*., *s*_1_ = 0.9994*z*_1_ − 0.0358*z*_2_ and *s*_2_ = 0.7193*z*_2_ − 0.6947*z*_1_ at *τ* = 4 ns. (Note that the effect of the PBCs on tICA CVs is not negligible for membrane permeation trajectories and thus tICA should be performed carefully as we discuss in the next section. For this analysis, we selected one of the trajectories whose tICA CVs were least affected by the PBCs.) Intriguingly, the first and second CVs represent the translational and orientational modes, respectively, as the ICs can be approximated as *s*_1_ ≈ *z*_1_ and *s*_2_ ≈ *z*_2_ − *z*_1_. The first CV carries the same information as *s*_0_ and *z*_CM_ because its value simply represents the vertical position of the COM of the TMB group. The second CV, which is the weighted difference between *z*_1_ and *z*_2_, is strongly correlated with the orientation angle *θ* of trimethoprim with respect to the surface normal of the membrane (*i*.*e*., the *z*-axis). This is clearly shown in Fig. S1 where the inner product of the directional vector *ŝ*_2_ of trimethoprim and the surface normal 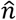 of the membrane shows a linear relationship with the value of *s*_2_: cos *θ* = −2.028*s*_2_ with *R*^2^ = 0.9989. Hence, *ŝ*_2_ is oriented towards the membrane center if *s*_2_ < 0 when *s*_1_ > 0 (near the upper leaflet) or if *s*_2_ > 0 when *s*_1_ < 0 (near the lower leaflet). The width of the curve simply reflects conformational fluctuations (particularly, elongation and contraction in the direction of *ŝ*_2_) of the drug molecule since the value of *s*_2_ will change significantly when 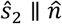(*i.e.*, cos*θ* = ±1) and minimally when 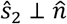(*i.e.*, cos*θ* = ±0) since both *z*_1_ and *z*_2_ are vertical positions with respect to the membrane COM.

**Figure 4.**
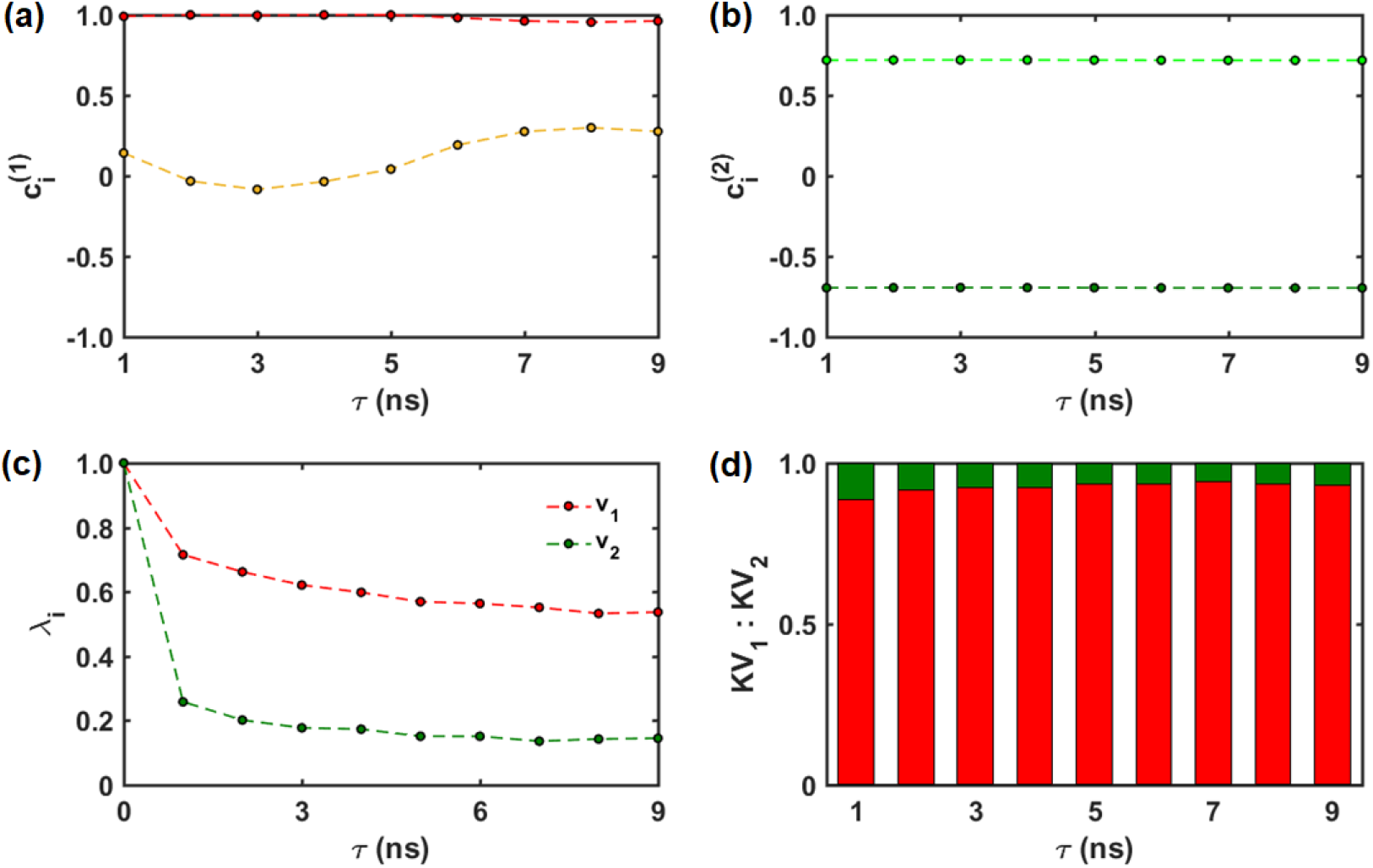
tICA results from one of the WTMetaD trajectories we obtained. (a) The first (red) and second (yellow) components 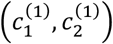 of the first eigenvector ***v***_1_, (b) the first (green) and second (chartreuse) components 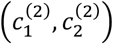 of the second eigenvector ***v***_2_, (c) the first (red, *i* = 1) and second (green, *i* = 2) eigenvalues, and (d) the ratio of the kinetic variances KV_*i*_ retained by the first (red, *i* = 1) and second (green, *i* = 2) eigenvectors as a function of the lag time *τ*.

Fig. 4(c) displays the eigenvalues *λ*_*i*_ corresponding to the first (red) and second (green) eigenvectors as a function of *τ*. A clear separation of the two timescales is detected. The eigenvalues quantify the largest autocorrelation functions of the projections of the molecular feature vector onto the slowest independent collective degrees of freedom at a given *τ* (Fig. S2). Physically, they are directly related to relaxation times 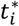 associated with the slow modes by

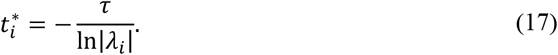

The dominant eigenvalue *λ*_1_ (and thus the slowest relaxation mode) is associated with the translational kinetics, and the associated relaxation time is 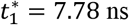 at *τ* = 4 ns. The other eigenvalue *λ*_2_ (and thus the second slowest relaxation mode) is associated with the orientational kinetics, and the associated relaxation time is 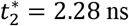 at *τ* = 4 ns. Fig. 4(d) shows the ratio of the kinetic variance KV_*i*_ retained by each IC at a given *τ* after dimensionality reduction, which is given by

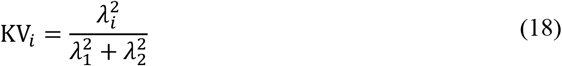

in this study. The kinetic variance is mostly retained by *s*_1_; for example, KV_1_ = 0.92 and KV_2_ = 0.08 at *τ* = 4 ns.

### 2D PMF along tICA CVs

Lastly, we should determine the quality of the slow CVs not only by their ability to distinguish different states but also by the recrossing frequency. The mean recrossing frequency (or the average rate of membrane crossings) is an important metric to determine the sampling efficiency of chosen CVs in membrane permeation simulations. Comparing recrossing frequencies serves as a reliable means to assess whether a crucial variable (slow mode) has been overlooked. It is widely known that if a slow mode is forgotten, the sampling process may suffer from hysteresis, leading to a lack of convergence in FES calculations^22, 61, 69^. We first computed the average 2D PMF spanned by our new CVs, *s*_1_ = 0.9964*z*_1_ − 0.0851*z*_2_ and *s*_2_ = 0.7197*z*_2_ − 0.6942*z*_1_, and identified the MFEP using the zero-temperature string method^62-63^ as shown in Fig. 5(a). The two metastable states exist at *A* = (−1.5, − 0.3) and *B* = (1.5, 0.3), and the transition state is located at (0, 0). Also, the PMF shows the preferential orientation of trimethoprim in the membrane; *ŝ*_2_ points away from the membrane center (*i*.*e*., the DAP group in the polar region and the TMB group in the hydrophobic region of the lipid bilayer) when the system is in the metastable states – see Fig. 3(a, c). The MFEP is a curve γ connecting critical points on an energy landscape *V* that satisfies (∇*V*)^⊥^(γ) = 0 and represents the most probable transition path in a large ensemble of the permeation processes.^70^ The MFEP starts with the DAP group oriented toward the lipid headgroups, while trimethoprim is in the aqueous region and above the lipid membrane (A in Fig. 5(a)), which is slightly favored over the opposite orientation (by ∼2.0 kJ/mol). From there, a barrier is encountered while TMB flips down into the hydrophobic region of phospholipids reversing the orientation of trimethoprim (B in Fig. 5(a)). The drug then crosses a larger barrier passing through the lipid tails and flipping again to orient DAP once again to the polar glycerol, phosphate, and headgroup region (C in Fig. 5(a)). The drug molecule then escapes from the lipid bilayer to the aqueous region with TMB flipping into the aqueous region (D in Fig. 5(a)). The 1D PMF along the MFEP is presented in Fig. 5(b). The heights of the small and large energy barriers are ∼17.9 kJ/mol and ∼47.1 kJ/mol, respectively, and the depth of the metastable states is ∼3.2 kJ/mol. The green points in Fig. 5(c) indicate the conformational space sampled in the 5 *μ*s-long unbiased simulation. Note that only the metastable states and the aqueous regions are sampled. Our visual inspection reveals one or two flipping events per 1 *μ*s on average at the region of lipid headgroups in the unbiased simulation. We analyzed the orientation of trimethoprim when it approaches the membrane surface from the aqueous region (Fig. S12(a)) and when it is buried in the membrane at the metastable states (marked with B and C in Fig. 5(a)) (Fig. S12(b)). This shows the probability distribution of the orientation angle *θ* of trimethoprim with respect to the surface normal of the nearest leaflet. We defined the orientation of the molecule as the vector connecting from the COM of the TMB group to the COM of the DAP group, or simply *ŝ*_2_. This confirms that the drug molecule reverses its orientation at the polar headgroup region such that TMB flips into the hydrophobic tail region.

**Figure 5.**
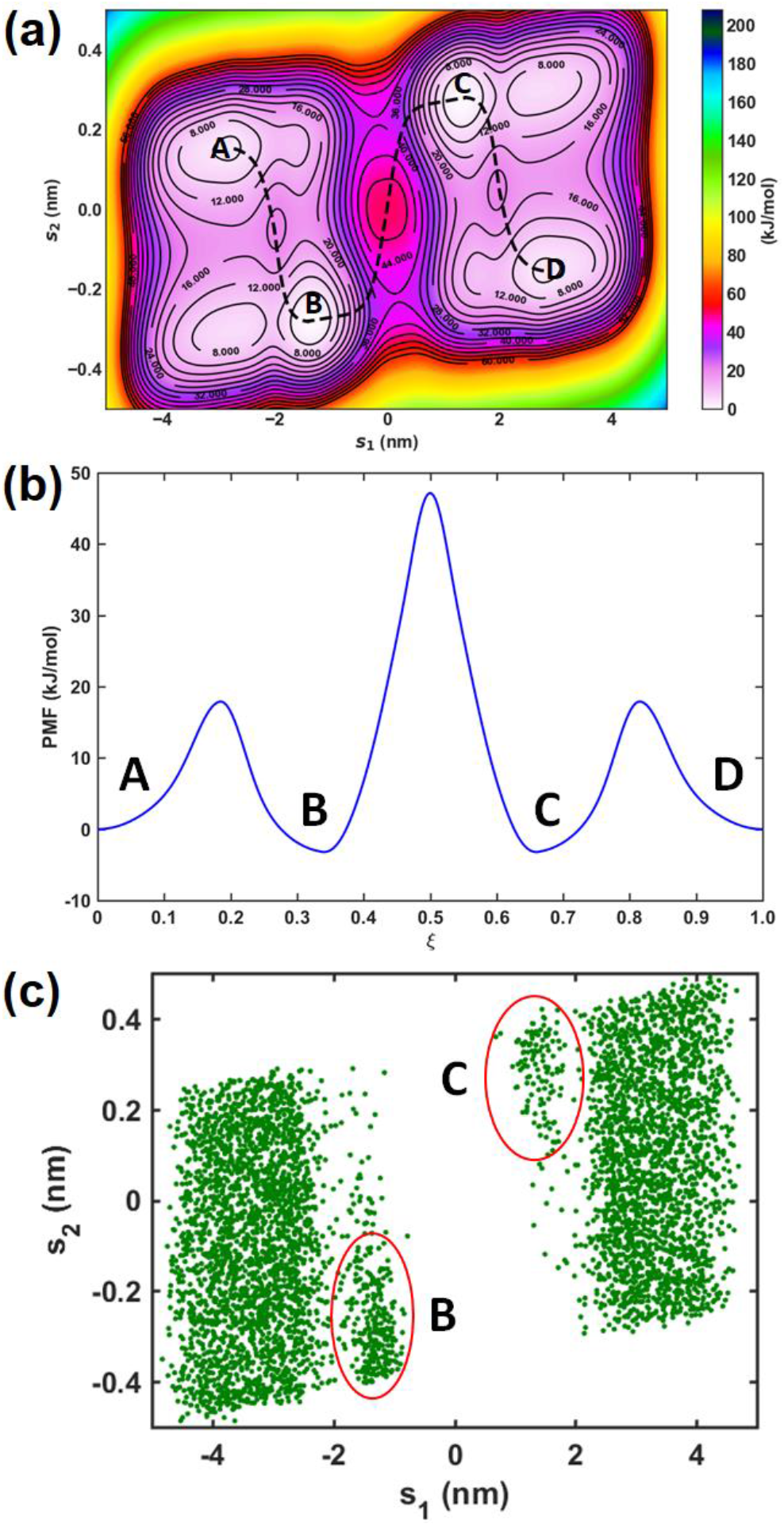
(a) Average 2D PMF along the tICA CVs *s*_1_ and *s*_2_. The dashed black line indicates the MFEP found from the zero-temperature string method. (b) 1D PMF along the MFEP ξ. The free energy in bulk water was set to zero. (c) Scatter plot showing all the (*s*_1_,*s*_2_) pairs sampled by the long-timescale standard MD simulation. B and C represent the metastable states.

To confirm our observation that the TMB group penetrates the lipid tails more readily, a series of analyses were also performed. First, tracking the z-coordinate of the COM of each group in the unbiased simulation (Fig. 6(a)) shows that when trimethoprim begins to enter the tail region (−2 nm < *Z*_COM_ < 2 nm), TMB goes deeper than DAP. Concurrently, the failed attempts to penetrate more frequently show DAP deeper, consistent with its preferential interaction with the lipid headgroups. We observed the same behavior in the biased simulations; TMB dives deeper each time trimethoprim attempts to penetrate and successful crossings are marked by a rapid flip (Fig. 6(b)).

**Figure 6.**
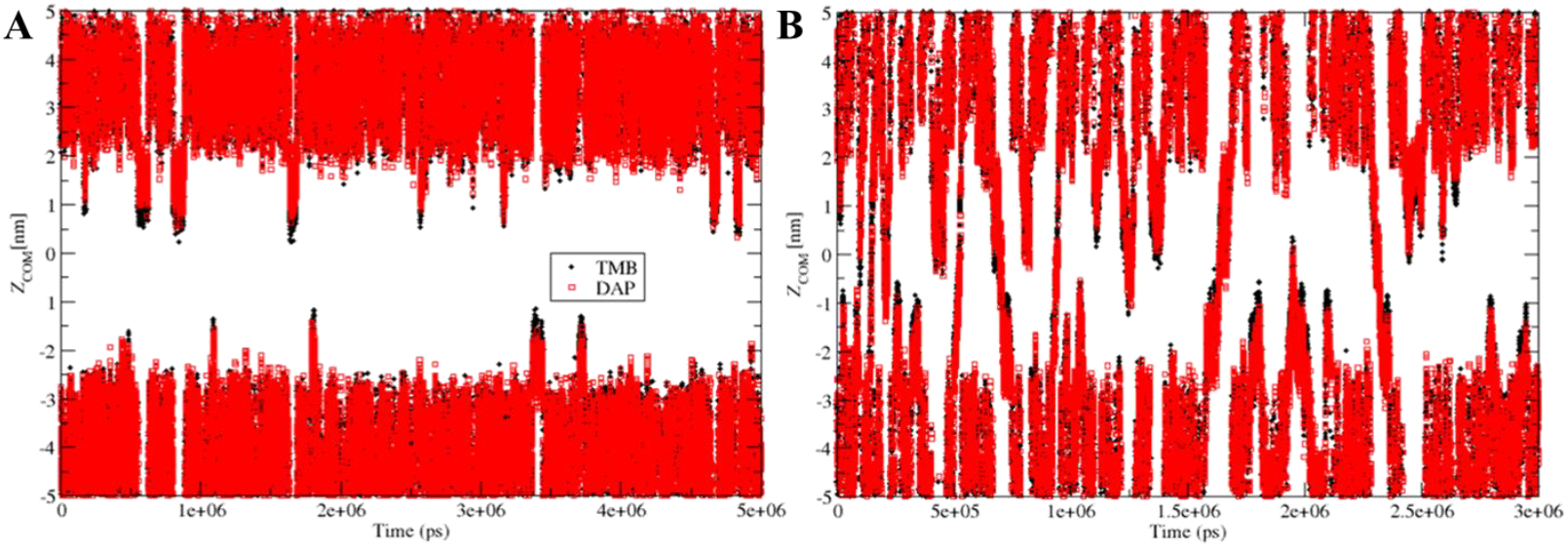
Time evolution of the *z*-coordinates of the COMs of DAP (red) and TMB (black) in the (a) unbiased and (b) biased simulations.

This information led to an investigation of which lipids the DAP and TMB groups interact with. Contact analysis was performed to explore different regions of the unbiased simulation system, focusing on membrane interactions. While both DAP and TMB have a similar number of contacts with the hydrophilic groups (*i*.*e*., choline and phosphate groups), TMB surpasses DAP in contacts with the deeper ester groups (Fig. S13(a-c)). This group is not only deeper in the membrane, but also includes the first carbons of the hydrophobic tails. The methoxy groups of TMB, due to their carbons having a slightly negative charge (*q* ∼ −0.1 *e*), facilitate burial in this region, where the carbons have a similar but positive charge (*q* ∼ 0.2 *e*). The oxygen atoms are also less negative (*q* ∼ −0.39 *e*) than the nitrogen atoms of the DAP group (*q* ∼ −0.75 *e*), thus having less electrostatic repulsion with ester oxygens. This trend (related to the carbons) is also observed in the contacts of each component with cholesterol, resulting in more contacts of TMB with POPC/cholesterol overall (Fig. S15 (a-b)). Finally, the contacts of TMB and DAP with the water molecules support the flip mechanism of trimethoprim when entering the membrane: when trimethoprim is in water, TMB has more total interactions with water molecules than DAP (equivalent when normalized per atom, Fig. S14(d)). When the molecule penetrates more deeply into the membrane (consistent with Fig. 6(a)), it flips directing the TMB portion into the more buried, hydrophobic tails, dropping water contacts to 0 in Fig. S13(d). Similar observations are apparent in the number of contacts per atom (Fig. S14).

The average interaction energy of DAP and TMB with each system component was also calculated to verify the described preferential interactions (Table S1). Based again on the unbiased simulation, TMB shows a larger average interaction energy than DAP with each component, simply because it is a larger molecule. But the relative increase is substantially larger for the ester group and cholesterol, and significantly reduced for the phosphate and choline groups.

Interestingly, our triple-flip mechanism is not consistent with the triple-flip model previously proposed by Sun *et al*.^*8*^ based on their TTMetaD simulations of trimethoprim permeating a homogeneous lipid bilayer composed only of POPC. In this model, the TMB group preferentially stays in the polar region of lipid headgroups, while the DAP group flips into the hydrophobic region of lipid tails due to its hydrophobicity. The mechanistic difference may be attributed to the effect of membrane cholesterol. However, there is no convincing evidence for the relative hydrophobicity for the two moieties. For example, Masoud *et al*.^71^ estimated the dipole moment of 2,4-diaminopyridimine to be ∼2.25-2.48 D based on their quantum-mechanical calculations, whereas the experimental dipole moment of 1,2,3-trimethoxybenzene was reported to be 2.25 D^72^. In addition, the TMB group possesses a benzene ring with only three hydrogen bond acceptors, whereas the DAP group contains a pyrimidine ring with two hydrogen bond donors and four hydrogen bond acceptors. Consequently, it is more energetically favorable for DAP to interact with the polar headgroups while TMB flips to water or the hydrophobic region of the lipid bilayer.

Mehdi *et al*.^73^ employed the State Predictive Information Bottleneck (SPIB) approach to investigate the permeation of a small benzoic acid (BA) molecule across a symmetric phospholipid bilayer, with the aim of identifying reaction coordinates (RCs) for enhanced sampling algorithms. In their study, the authors considered two pivotal order parameters (OPs) pertaining to BA permeation, namely the membrane-BA *z*-distance (*d*_1*Z*_) and the angle formed between BA and the *z*-axis (θ_*Z*_). Analysis of the SPIB RCs identifies key steps involved in the entry and exit mechanisms of BA. Prior to penetrating the membrane, the polar region of BA initiates interactions with the membrane surface, acting as an anchor. This initial interaction facilitates the inward flipping of the BA benzene ring, enabling it to enter the membrane. Once inside, the polar region of BA continues to engage with the headgroup regions of the membrane, while the hydrophobic benzene ring of BA orients itself close to the center of the membrane, establishing interactions with the lipid tails. Ultimately, after surmounting the central barrier, which corresponds to the flipping of BA within the membrane, BA translocates to the opposite leaflet of the membrane, initiating interactions between the polar region of BA and the headgroup of the membrane. Remarkably, this mechanism bears a striking resemblance to the triple-flip model observed in the permeation process of trimethoprim.

The mean recrossing frequency (or the average rate of membrane crossings) is an important metric to determine the sampling efficiency of chosen CVs in membrane permeation simulations. Fig. S3 displays how *z*_1_ changes over time when (a) *s*_0_ and (b) *s*_1_ and *s*_2_ are biased in biased trajectories. We found that the mean recrossing frequency of *s*_0_, 3.3 ± 1.3 *μs*^−1^, increases to 5.0 ± 2.2 *μs*^−1^ (∼1.5 times increase) when *s*_1_ and *s*_2_ are used as CVs. This is only a modest increase in efficiency, likely due to the translational position captured by both *s*_0_ and *s*_1_being the dominant slow mode.

Lastly, we found that the same tICA-MetaD procedure cannot be applied to TTMetaD trajectories. TTMetaD was developed to prevent undesirable situations that may arise from the inappropriate choice of the bias factor which controls the speed at which the Gaussian height decreases. TTMetaD aggressively tempers the height of Gaussian hills only after basins, whose locations are roughly defined prior to simulations, are relatively full. We generated one 1 *μs*-long TTMetaD trajectory with the same parameters as in our previous study^47^ and plotted how rescaled *z*_1_, *V*(*s,t*), *c*(*t*), and *γ*(*t*) evolve with time in Fig. S4. We observed that the difference between *V*(*s,t*) and *c*(*t*) is extremely large in TTMetaD compared to WTMetaD; *γ*(*t*) fluctuates mostly between 10 and −50 kJ/mol in Fig. 2(d) and around −160 kJ/mol in the lower right panel of Fig. S4. Therefore, the time step becomes extremely small when rescaled according to Eq. 14, and consequently, tICA cannot be performed for a reasonable range of *τ*.

### Effect of the Periodic Boundary Conditions on the Slow CVs

A significant problem appears when the time series analysis is done directly on the data points obtained from membrane permeation simulations because the PBCs make the behavior of the permeant “unphysical” when it crosses the boundaries (particularly, in the *z*-dimension for our systems). The tICA algorithm would recognize the boundary conditions as the unrealistic transfer (or “teleportation”) of the molecule from one end of the simulation box to the other without traversing the physical space between them. We performed tICA on 10 different reweighted trajectories to compute the slow CVs at 10 different lag times ranging from 1 to 10 ns (see Fig. 7(a) and Fig. S5). We observed that the first eigenvectors are scattered on the domain defined by 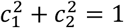 and −π *≤ θ ≤ π*/2 where *θ* is the angle between the positive *c*_1_-axis and the vector. The data points lie on the circumference of a unit circle as the eigenvectors are normalized. Interestingly, the second eigenvectors are slightly scattered around (−0.7,0.7) and show a high level of precision compared to the first eigenvectors since the orientational change of trimethoprim is less affected by the PBCs than its translational motion. To identify the center of each eigenvector in the data, we used the Partitioning Around Medoids (PAM) clustering algorithm, which is similar to *k*-means clustering but requires the centroid of each cluster to be one of the input data points. We found that the centers of the first and second eigenvectors are (0.639,−0.769) and (−0.707,0.707), respectively. Thus, when the PBCs are applied, the translational mode is not well captured by tICA as the drug molecule can reach the other side of the membrane rapidly without crossing the membrane.

**Figure 7.**
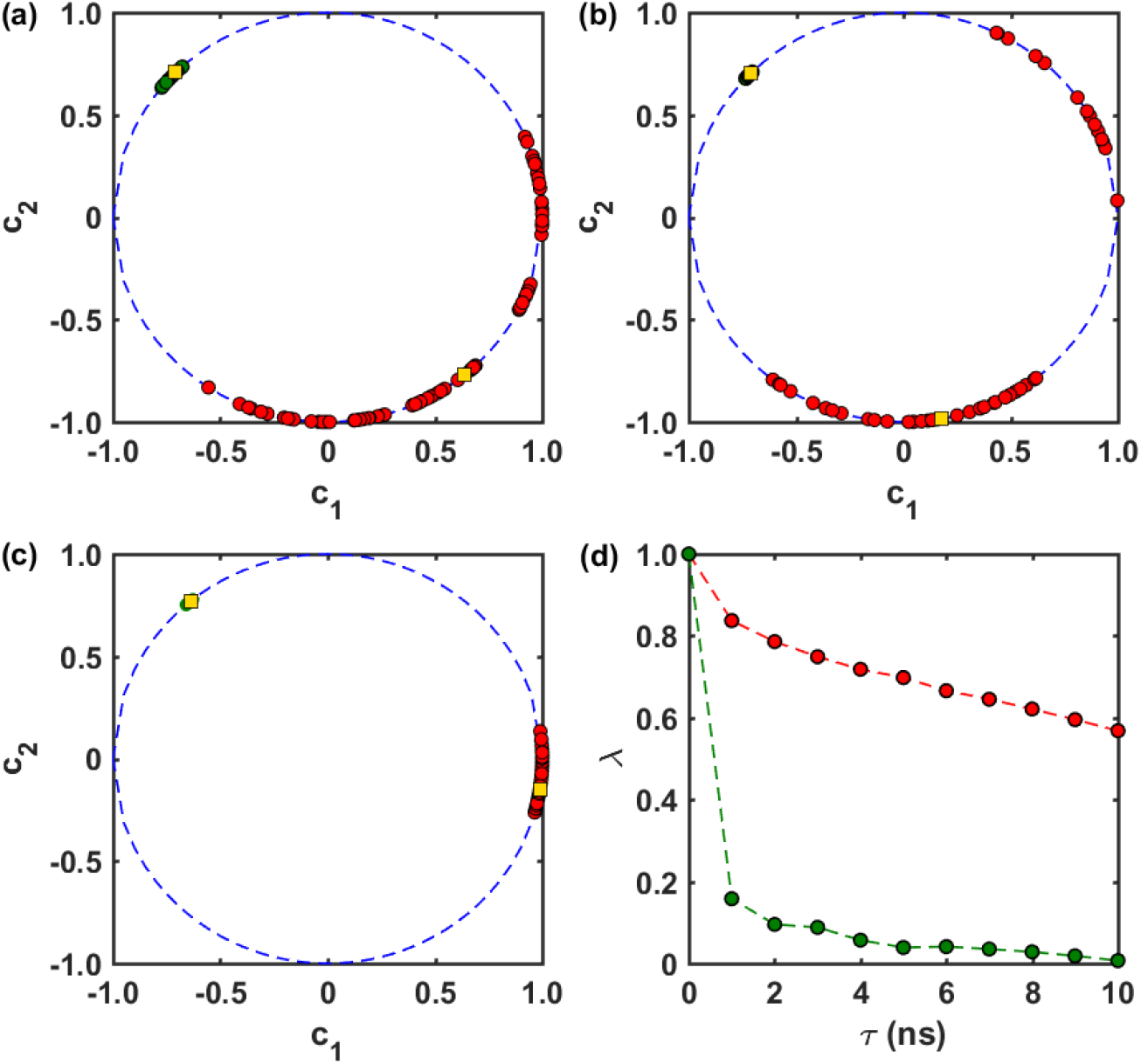
The first (red) and second (green) eigenvectors (*c*_1_,*c*_2_) obtained from tICA-MetaD with (a) the PBCs, (b) the upper and lower walls, and (c) the absolute values of the molecular features. The yellow squares represent the medoids. (d) The first (red) and second (green) eigenvalues as a function of *τ* obtained from tICA on one of the full biased trajectories.

We examined two different approaches to address the PBC issue. In the first approach, we applied harmonic potentials to construct lower and upper walls in the water region below and above the bilayer and thus trap trimethoprim inside the space between the two boundaries in the *z*-direction. The free energy surface along *z*_1_ and *z*_2_ in the presence of the walls is shown in Fig. S6, and we did not see any significant difference from the one obtained in the absence of the walls in previous works.^8, 47^ We applied tICA to 5 different reweighted trajectories to calculate the slow CVs at 10 different lag times ranging from 1 to 10 ns (see Fig. 7(b) and Fig. S7). As in the case of the PBCs, the first eigenvectors are scattered on the circumference of a unit circle within −π *≤ θ ≤ π*/2 but characterized by a higher degree of dispersion. This is likely because the behavior of the drug molecule is unnatural due to its collisions with the walls in the aqueous region. However, the second eigenvectors capture the orientational mode more precisely compared to the case in which the walls are absent since boundary crossings are physically prevented by the walls. Our *k*-medoids analysis reveals that the centroids are (0.179,−0.984) and (−0.712,0.702) for the first and second eigenvectors, respectively. The centroid of the first eigenvectors still captures the translational motion of the drug molecule to a considerable extent with the coefficient of *z*_2_ much larger in magnitude than that of *z*_1_, but their level of dispersion does not make it ideal to be selected as the translational CV.

In the second approach, we took absolute values of the molecular features *z*_1_ and *z*_2_ in the time series data to make them translationally invariant. The molecular features are any real numbers in the range of [−*d*_*z*_/2,*d*_*z*_/2] where *d*_*z*_ is the length of the simulation box in the *z*-direction. Their absolute values lose the information about which leaflet of the membrane is closer to trimethoprim but only retain the information about how far the drug molecule is away from the membrane as they can take any real numbers in the range of [0,*d*_*z*_/2]. The tICA results are summarized in Fig. 7(c) and Fig. S8. Surprisingly, the eigenvectors are highly stable with respect to lag times and consistent over different trajectories. Furthermore, the levels of dispersion for both the first and second eigenvectors are significantly reduced. The medoids are (0.989,−0.148) and (−0.638,0.770) for the first and second eigenvectors, respectively. Therefore, the second approach achieves the highest level of precision for both the first and second eigenvectors and the highest level of accuracy for the first eigenvector *but does not predict well the orientational mode of the drug molecule*. However, the sampling efficiency was not improved when we used the absolute values either as direct CVs or in the definitions of *s*_1_ and *s*_2_. The former is expected since |*z*_1_| and |*z*_2_| separately do not distinguish the upper and lower regions of the lipid bilayer (upper when *z*_1_,*z*_2_ *>* 0, and lower when *z*_1_,*z*_2_ < 0); taking the absolute values removes the locational information and thus biasing them cannot induce trimethoprim to cross the membrane. When we biased 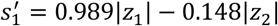 and 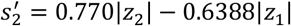, the mean recrossing frequency over three replicas was only 1.3 ± 0.5 *μs*^−1^ (Fig. S9). Thus, taking the absolute values extracts the coefficients of the translational CV in a more reliable manner but it poorly describes the orientational motion of the drug molecule. Also, using the absolute values in the definitions of CVs does not necessarily improve sampling performance of WTMetaD due to a loss of the locational information of the drug molecule.

### Effect of Data Reweighting

Data reweighting recovers information about the unbiased dynamics of the system using the conformations collected from biased simulations and their statistical weights. However, fast and accurate convergence to the underlying unbiased free energy is not always guaranteed depending on the reweighting procedure, the enhanced sampling technique, and the stage of the simulation.^45^ In our previous work,^47^ we obtained tICA CVs directly from short biased trajectories that involve at least one permeation event without data reweighting. Initially, a TTMetaD simulation lasting 70 ns was performed, utilizing *z*_1_ and *z*_2_ as CVs to generate an initial trajectory involving at least one membrane crossing. Subsequently, the *z*-positions of five heavy atoms in the drug molecule, namely *Z*_1_ to *Z*_5_, were calculated relative to the COM of the lipid bilayer. tICA was then applied to this trajectory to identify the most dominant eigenvector, corresponding to the slowest mode. Another TTMetaD simulation was conducted, incorporating the tICA CV and the z-position of the COM of the drug molecule (denoted as *Z*). Comparisons between the results obtained from simulations utilizing tICA-based CVs and the traditional CVs (*z*_1_ and *z*_2_) revealed that the tICA CVs achieved faster convergence and yielded more accurate outcomes. Essentially, the use of tICA CVs enhances the efficiency of potential of mean force (PMF) calculations, while concurrently providing additional insights into the permeation mechanism. Specifically, the tICA CV unveiled a subtle influence of cholesterol on the resistance of the lipid head group region to permeation, which was not observed when employing the canonical CVs. It is, therefore, necessary to understand the effect of data reweighting on tICA CVs and determine the minimum number of rare events required for tICA CVs to converge. This information can be used to further optimize the tICA-MetaD procedure for future membrane permeation simulations.

We first extracted from our WTMetaD trajectories the subtrajectories (or segments of trajectories of specified duration) that contain only one to seven membrane crossings, and grouped them into seven sets of 10 subtrajectories according to the number of membrane crossings included (see Fig. S10 for an example of each set). We then performed tICA on each subtrajectory with and without data reweighting. We chose |*z*_1_| and |*z*_2_| as the molecular features for tICA because the eigenvectors show the highest degree of convergence when the absolute values are used (Fig. 7(c)). Fig. 8 summarizes tICA results with data reweighting. Here, we adopted the same color scheme as in Fig. 4 and displayed in order the 100 eigenvector pairs we obtained for each set (since the analysis was performed at 10 different lag times on 10 subtrajectories) to evaluate their converging behavior. We also included the results from the 10 full biased trajectories (Fig. 8(h)) for the purpose of comparison. We observed that tICA does not capture the slow modes properly when subtrajectories contain only one permeation event because *τ* is relatively large for the length of the subtrajectories. We found that at least five permeation events are needed for the tICA CVs to achieve a high level of convergence (Fig. 8(e)). The medoids are (0.996,−0.087) and (−0.635,0.772) for the first and second eigenvectors, respectively, in Fig. 8(e), and (0.989,−0.148) and (−0.638,0.770) for the full trajectories in Fig. 8(h). The large fluctuations of the second component of the first eigenvector (yellow curves) arise mostly from the effect of vector normalization. For example, considering vectors only in the first quadrant, when the first component changes from 1 to 0.99, the second component changes from 0 to 0.14; when the first component changes from 0.70 to 0.69, the second component only changes from 0.71 to 0.72.

**Figure 8.**
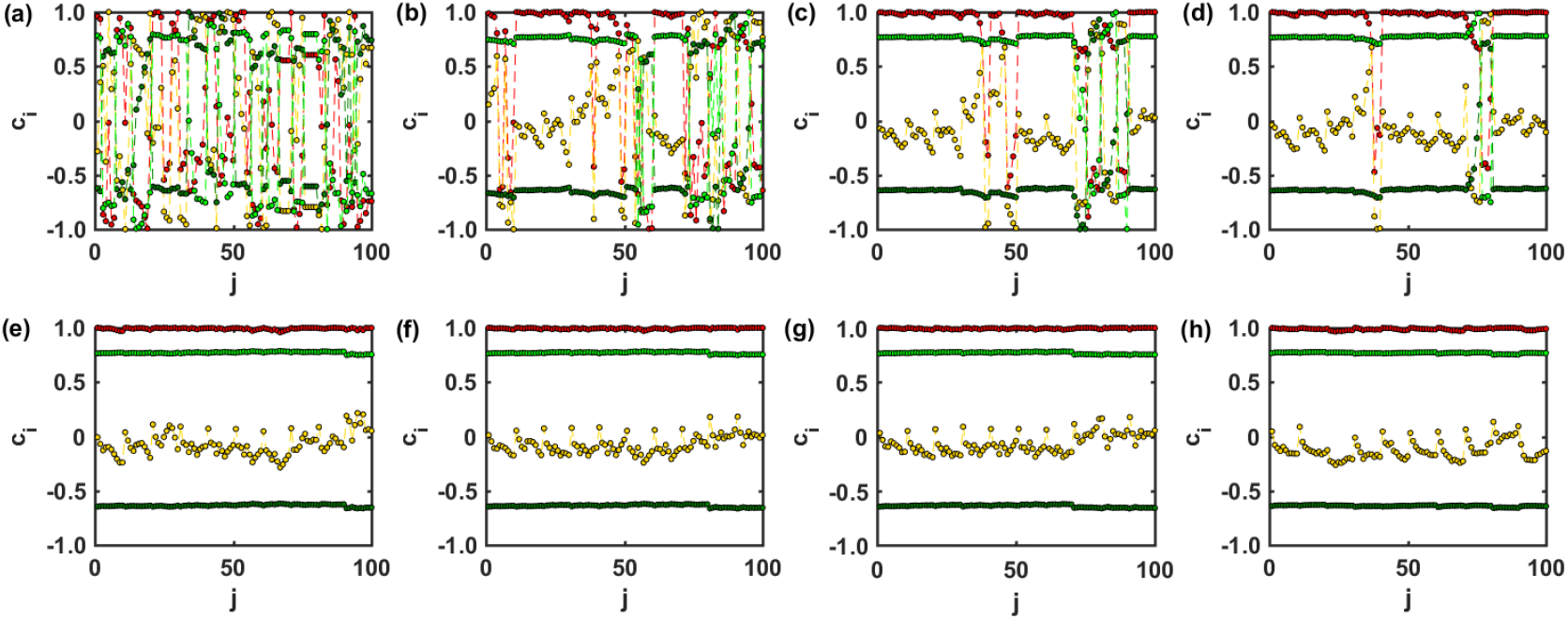
Results from tICA on reweighted subtrajectories involving (a) one, (b) two, (c) three, (d) four, (e) five, (f) six, and (g) seven membrane crossings and (h) on the full biased trajectories after reweighting. The same color scheme was used as in Fig. 4(a-b) to represent the components of the first and second eigenvectors.

Next, we omitted data reweighting and applied tICA directly to each subtrajectory and full biased trajectories (Fig. 9). The eigenvectors are well converged when at least five membrane crossings are involved. The medoids are (0.968,−0.250) and (−0.678,0.735) for the first and second eigenvectors, respectively, in Fig. 9(e), and (0.965,−0.260) and (−0.676,0.737) for the full trajectories in Fig. 9(h). However, the eigenvectors are quite different than those obtained from the reweighted trajectories. The first eigenvectors deviate more from the coefficients of the translational CV (*s*_1_ ≈ *z*_1_) whereas the second eigenvectors get slightly closer to the coefficients of the orientational CV (*s*_2_ ≈ 0.7*z*_2_ − 0.7*z*_1_) upon removal of data reweighting. We also observed that the orientational mode is not significantly affected by data reweighting even when the absolute values of the molecular features are not taken and the PBCs are still present, as clearly seen in Fig. S5 (dark and light lines for the first and second components of the second eigenvector, respectively); the second eigenvectors are highly consistent and converge to *s*_2_ even without reweighting. These observations can be attributed to the fact that the original bias was applied to the suboptimal CV *s*_0_ simply to increase the translational kinetics and thus the rate of membrane crossing of trimethoprim. In addition, since the orientational kinetics is much faster than the translational kinetics, we may assume that the two modes are not strongly coupled to each other. Consequently, without data reweighting the Boltzmann statistics of the translational mode cannot be recovered, whereas the orientational CV can be obtained with an acceptable degree of accuracy and convergence.

**Figure 9.**
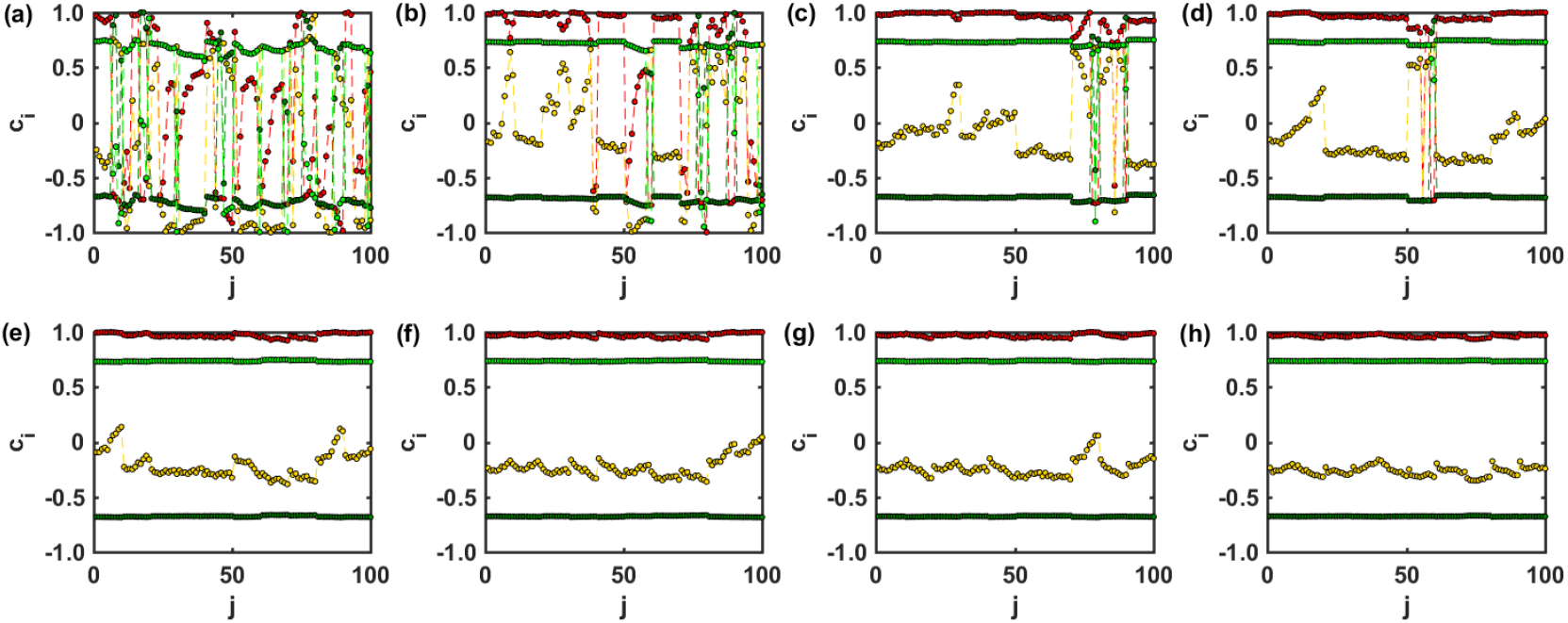
Results from tICA on subtrajectories involving (a) one, (b) two, (c) three, (d) four, (e) five, (f) six, and (g) seven membrane crossings and (h) on the full biased trajectories without data reweighting. The same color scheme was used as in Fig. 8 to represent the components of the first and second eigenvectors.

## CONCLUSION

In this work, we applied the tICA-MetaD procedure proposed by McCarty and Parrinello^41^ to trimethoprim membrane permeation to better understand the effectiveness, limitations, and best-practices of this methodology for membrane permeation simulations. Specifically, we sought to identify the slowest collective degrees of freedom and see if they improved simulation efficiency while also investigating the effects of the PBCs, the number of rare events in the original biased simulations, and data reweighting on the accuracy and convergence of tICA CVs. We found that the same tICA-MetaD scheme cannot be applied to TTMetaD trajectories due to extremely small reweighting factors and rescaled time steps. We observed that tICA-MetaD captures the translational and orientational modes separately, and the use of the tICA CVs accelerates the convergence of WTMetaD PMF calculations compared to the suboptimal CV initially selected. However, the PBCs may be detrimental to tICA in membrane permeation trajectories, particularly for identification of the translational CV. Adding harmonic restraints to prevent PBC crossing does not solve the problem because it causes unnatural, rapid diffusive behavior of the drug molecule in the aqueous region, whereas taking absolute values of the molecular features during tICA analysis can reliably recover the translational CV. In contrast, the orientational CV is not significantly affected by the PBCs and the walls, but is also not well captured when the absolute values are taken. Thus, taking absolute values is only useful for extracting the translational CV, and a method that captures both modes without PBC artefacts would be an important contribution to the field. Interestingly, we found that only five permeation events are sufficient for the tICA CVs to achieve convergence regardless of data reweighting, but the first eigenvector is quite different from the translational CV when data reweighting is omitted. Based on our results, we suggest that (1) the tICA-MetaD procedure can be applied to a short initial WTMetaD trajectory (after the transient time) that involves at least five membrane crossings, and (2) absolute values of the molecular features should be used along with data reweighting for correction of the translational CV.

It is indeed a non-trivial task to recover slow modes of the unbiased system through reweighting of the biased trajectory. Bonati *et al*.^74^ extended the approach initially proposed by McCarty and Parrinello^41^ in two significant ways. Firstly, they introduced a nonlinear variant of VAC which offers enhanced variational flexibility. This is achieved by employing neural networks to learn basis functions through a nonlinear transformation of descriptors for the variational principle. Second, they proposed novel strategies for collecting initial trajectories, such as sampling generalized ensembles instead of relying on trial CVs, as well as for fully using information gathered during the initial trajectory. Additionally, they utilize on-the-fly probability-enhanced sampling (OPES) to construct the bias potential, which presents several advantages over metadynamics and other methods. The deep-tICA method developed by Bonati *et al*.^74^ allows one to analyze a biased simulation trajectory, identify slow modes that impede convergence, and subsequently, accelerate them.

Chen and Chipot^75^ conducted an in-depth investigation into the use of classical autoencoders (AEs), time-lagged AEs (TAEs), modified TAEs, VAMPnets, and state-free reversible VAMPnets (SRVs) for deep learning-based CV discovery in molecular processes. Their findings revealed that classical AEs capture high-variance modes instead of slow modes, and this limitation can be overcome by turning to the time-series-based models. To determine an appropriate reweighting strategy for iterative learning, they compared the accuracy of Koopman reweighting^34^ and reweighting by uneven time intervals^67^. Their results demonstrated that with a suitable reweighting method, SRVs coupled with iterative learning facilitate the discovery of CVs^75^.

Additionally, there are alternative reweighting schemes that have been proposed in the literature. For instance, Girsanov reweighting is a technique designed to correct the probability weights of dynamical pathways gathered under a perturbed Hamiltonian to those collected under an unperturbed Hamiltonian^76-77^. This is accomplished by dynamically reweighting the transition density elements along the path through phase space^76-77^. Donati and Keller^77^ introduced a methodology that applies the Girsanov reweighting scheme to metadynamics simulations, allowing for the recovery of accurate dynamic properties of a molecular system from an enhanced sampling simulation. Another noteworthy approach is the Girsanov Reweighting Enhanced Sampling Technique (GREST) proposed by Shmilovich and Ferguson^76^. GREST is an adaptive sampling scheme that alternates rounds of data-driven slow CV discovery and enhanced sampling along these coordinates. In their work, the authors employed state-free reversible VAMPnets (SRVs) for the identification of slow CVs, while the Girsanov formalism facilitated the estimation of dynamical observables under an unbiased Hamiltonian from biased trajectories. This formalism provides guidelines for reweighting trajectories in the biased path ensemble according to both thermodynamic and integrator-specific dynamical path weights^76^.

Future work should investigate how to select a minimal set of molecular features for optimal CVs, how to incorporate deep learning algorithms to add flexibility in feature selection and evade the linearity problem of tICA, and how the results may change when different reweighting algorithms are used for WTMetaD and TTMetaD.

## ASSOCIATED CONTENT

### Supporting Information

The following files are available free of charge.

Figures S1-S16 and Table S1 (PDF)

## AUTHOR INFORMATION

### Author Contributions

MO and JMJS designed the research. GCAH and MO carried out the simulations and analyses. All authors interpreted the results and wrote the manuscript.

## ACKNOWLEDGMENT

The authors gratefully acknowledge support from NIH NIGMS (R35GM143117) and computational support from EXPANSE at the San Diego Supercomputing Center through the ACCESS program (allocation MCB200018) supported by NSF (grants #2138259, #2138286, #2138307, #2137603, and #2138296), as well as the Center for High Performance Computing at the University of Utah.

## ABBREVIATIONS

COM: center of mass
CGenFF: CHARMM general force field
CV: collective variable
DAP: diaminopyrimidine
IC: independent component
ISD: inhomogeneous solubility-diffusion
KV: kinetic variance
LINCS: linear constraint solver
MetaD: metadynamics
MFEP: minimum free energy path
MD: molecular dynamics
PBC: periodic boundary condition
PO4: phosphate
POPC: phosphatidylcholine
PMF: potential of mean force
PC: principal component
PCA: principal component analysis
SPME: smooth particle-mesh Ewald
tICA: time-lagged independent component analysis
TTMetaD: transition-tempered metadynamics
TMB: trimethoxybenzyl
VMD: visual molecular dynamics
WTMetaD: well-tempered metadynamics

## REFERENCES

1. Buchenberg, S.; Schaudinnus, N.; Gerhard, S., Hierarchical Biomolecular Dynamics: Picosecond Hydrogen Bonding Regulates Microsecond Conformational Transitions. J. Chem. Theory Comput. 2015, 11 (3), 1330–1336.

2. Henzler-Wildman, K.; Kern, D., Dynamic personalities of proteins. Nature 2007, 450 (7172), 964–972.

3. Lewandowski, J. R.; Halse, M. E.; Blackledge, M.; Emsley, L., Direct observation of hierarchical protein dynamics. Science 2015, 348 (6234), 578–581.

4. Chen, M., Collective variable-based enhanced sampling and machine learning. Eur. Phys. J. B 2021, 94 (10), 211.

5. Ghaemi, Z.; Minozzi, M.; Carloni, P.; Laio, A., A Novel Approach to the Investigation of Passive Molecular Permeation through Lipid Bilayers from Atomistic Simulations. J. Phys. Chem. B 2012, 116 (29), 8714–8721.

6. Awoonor-Williams, E.; Rowley, C. N., Molecular simulation of nonfacilitated membrane permeation. Biochim. Biophys. Acta - Biomembr. 2016, 1858 (7, Part B), 1672–1687.

7. Sugita, M.; Sugiyama, S.; Fujie, T.; Yoshikawa, Y.; Yanagisawa, K.; Ohue, M.; Akiyama, Y., Large-Scale Membrane Permeability Prediction of Cyclic Peptides Crossing a Lipid Bilayer Based on Enhanced Sampling Molecular Dynamics Simulations. J. Chem. Inf. Model. 2021, 61 (7), 3681–3695.

8. Sun, R.; Dama, J. F.; Tan, J. S.; Rose, J. P.; Voth, G. A., Transition-Tempered Metadynamics Is a Promising Tool for Studying the Permeation of Drug-like Molecules through Membranes. J. Chem. Theory Comput. 2016, 12 (10), 5157–5169.

9. Pokhrel, N.; Maibaum, L., Free Energy Calculations of Membrane Permeation: Challenges Due to Strong Headgroup–Solute Interactions. J. Chem. Theory Comput. 2018, 14 (3), 1762–1771.

10. Krämer, A.; Ghysels, A.; Wang, E.; Venable, R. M.; Klauda, J. B.; Brooks, B. R.; Pastor, R. W., Membrane permeability of small molecules from unbiased molecular dynamics simulations. J. Chem. Phys. 2020, 153 (12), 124107.

11. Marrink, S.-J.; Berendsen, H. J. C., Simulation of water transport through a lipid membrane. J. Phys. Chem. 1994, 98 (15), 4155–4168.

12. Venable, R. M.; Krämer, A.; Pastor, R. W., Molecular Dynamics Simulations of Membrane Permeability. Chem. Rev. 2019, 119 (9), 5954–5997.

13. Davoudi, S.; Ghysels, A., Sampling efficiency of the counting method for permeability calculations estimated with the inhomogeneous solubility–diffusion model. J. Chem. Phys. 2021, 154 (5).

14. Vervust, W.; Zhang, D. T.; van Erp, T. S.; Ghysels, A., Path sampling with memory reduction and replica exchange to reach long permeation timescales. Biophys. J.

15. Swenson, D. W. H.; Prinz, J.-H.; Noe, F.; Chodera, J. D.; Bolhuis, P. G., OpenPathSampling: A Python Framework for Path Sampling Simulations. 1. Basics. J. Chem. Theory Comput. 2019, 15 (2), 813–836.

16. Yang, Y. I.; Shao, Q.; Zhang, J.; Yang, L.; Gao, Y. Q., Enhanced sampling in molecular dynamics. J. Chem. Phys. 2019, 151 (7), 070902.

17. Ghysels, A.; Roet, S.; Davoudi, S.; van Erp, T. S., Exact non-Markovian permeability from rare event simulations. Phys. Rev. Res. 2021, 3 (3), 033068.

18. Kaptan, S.; Vattulainen, I., Machine learning in the analysis of biomolecular simulations. Adv. Phys.: X 2022, 7 (1), 2006080.

19. Bussi, G.; Laio, A., Using metadynamics to explore complex free-energy landscapes. Nat. Rev. Phys. 2020, 2 (4), 200–212.

20. Trapl, D.; Horvacanin, I.; Mareska, V.; Ozcelik, F.; Unal, G.; Spiwok, V., Anncolvar: Approximation of Complex Collective Variables by Artificial Neural Networks for Analysis and Biasing of Molecular Simulations. Front. Mol. Biosci. 2019, 6.

21. Sidky, H.; Chen, W.; Ferguson, A. L., Machine learning for collective variable discovery and enhanced sampling in biomolecular simulation. Mol. Phys. 2020, 118 (5), e1737742.

22. Valsson, O.; Tiwary, P.; Parrinello, M., Enhancing Important Fluctuations: Rare Events and Metadynamics from a Conceptual Viewpoint. Annu. Rev. Phys. Chem. 2016, 67 (1), 159–184.

23. Bernetti, M.; Bertazzo, M.; Masetti, M., Data-Driven Molecular Dynamics: A Multifaceted Challenge. Pharmaceuticals 2020, 13 (9), 253.

24. Glielmo, A.; Husic, B. E.; Rodriguez, A.; Clementi, C.; Noé, F.; Laio, A., Unsupervised Learning Methods for Molecular Simulation Data. Chem. Rev. 2021, 121 (16), 9722–9758.

25. Lange, O. F.; Grubmüller, H., Can Principal Components Yield a Dimension Reduced Description of Protein Dynamics on Long Time Scales? J. Phys. Chem. B 2006, 110 (45), 22842–22852.

26. Sittel, F.; Jain, A.; Stock, G., Principal component analysis of molecular dynamics: On the use of Cartesian vs. internal coordinates. J. Chem. Phys. 2014, 141 (1), 014111.

27. Amadei, A.; Linssen, A. B. M.; de Groot, B. L.; van Aalten, D. M. F.; Berendsen, H. J. C., An Efficient Method for Sampling the Essential Subspace of Proteins. J. Biomol. Struct. Dyn. 1996, 13 (4), 615–625.

28. Amadei, A.; Linssen, A. B. M.; Berendsen, H. J. C., Essential dynamics of proteins. Proteins 1993, 17 (4), 412–425.

29. Daidone, I.; Amadei, A., Essential dynamics: foundation and applications. Wiley Interdiscip. Rev. Comput. Mol. Sci. 2012, 2 (5), 762–770.

30. David, C. C.; Jacobs, D. J., Principal Component Analysis: A Method for Determining the Essential Dynamics of Proteins. In Protein Dynamics: Methods and Protocols, Livesay, D. R., Ed. Humana Press: Totowa, NJ, 2014; pp 193–226.

31. Naritomi, Y.; Fuchigami, S., Slow dynamics of a protein backbone in molecular dynamics simulation revealed by time-structure based independent component analysis. J. Chem. Phys. 2013, 139 (21), 215102.

32. Molgedey, L.; Schuster, H. G., Separation of a mixture of independent signals using time delayed correlations. Phys. Rev. Lett. 1994, 72 (23), 3634–3637.

33. Noé, F.; Clementi, C., Collective variables for the study of long-time kinetics from molecular trajectories: theory and methods. Curr. Opin. Struct. Biol. 2017, 43, 141–147.

34. Wu, H.; Nüske, F.; Paul, F.; Klus, S.; Koltai, P.; Noé, F., Variational Koopman models: Slow collective variables and molecular kinetics from short off-equilibrium simulations. J. Chem. Phys. 2017, 146 (15), 154104.

35. Schultze, S.; Grubmüller, H., Time-Lagged Independent Component Analysis of Random Walks and Protein Dynamics. J. Chem. Theory Comput. 2021, 17 (9), 5766–5776.

36. Pérez-Hernández, G.; Paul, F.; Giorgino, T.; Fabritiis, G. D.; Noé, F., Identification of slow molecular order parameters for Markov model construction. J. Chem. Phys. 2013, 139 (1), 015102.

37. Sittel, F.; Stock, G., Perspective: Identification of collective variables and metastable states of protein dynamics. J. Chem. Phys. 2018, 149 (15), 150901.

38. Nüske, F.; Keller, B. G.; Pérez-Hernández, G.; Mey, A. S. J. S.; Noé, F., Variational Approach to Molecular Kinetics. J. Chem. Theory Comput. 2014, 10 (4), 1739–1752.

39. Noé, F.; Nüske, F., A Variational Approach to Modeling Slow Processes in Stochastic Dynamical Systems. Multiscale Model. Simul. 2013, 11 (2), 635–655.

40. M. Sultan, M.; Pande V. S., tICA-Metadynamics: Accelerating Metadynamics by Using Kinetically Selected Collective Variables. J. Chem. Theory Comput. 2017, 13 (6), 2440–2447.

41. McCarty, J.; Parrinello, M., A variational conformational dynamics approach to the selection of collective variables in metadynamics. J. Chem. Phys. 2017, 147 (20), 204109.

42. Brotzakis, Z. F.; Parrinello, M., Enhanced Sampling of Protein Conformational Transitions via Dynamically Optimized Collective Variables. J. Chem. Theory Comput. 2019, 15 (2), 1393–1398.

43. Zhang, Y.-Y.; Niu, H.; Piccini, G.; Mendels, D.; Parrinello, M., Improving collective variables: The case of crystallization. J. Chem. Phys. 2019, 150 (9), 094509.

44. Dama, J. F.; Rotskoff, G.; Parrinello, M.; Voth, G. A., Transition-Tempered Metadynamics: Robust, Convergent Metadynamics via On-the-Fly Transition Barrier Estimation. J. Chem. Theory Comput. 2014, 10 (9), 3626–3633.

45. Schäfer, T. M.; Settanni, G., Data Reweighting in Metadynamics Simulations. J. Chem. Theory Comput. 2020, 16 (4), 2042–2052.

46. Crellin, E.; Mansfield, K. E.; Leyrat, C.; Nitsch, D.; Douglas, I. J.; Root, A.; Williamson, E.; Smeeth, L.; Tomlinson, L. A., Trimethoprim use for urinary tract infection and risk of adverse outcomes in older patients: cohort study. Br. Med. J. 2018, 360, k341.

47. Aydin, F.; Durumeric, A. E. P.; Hora, G. C. A. d.; Nguyen, J. D. M.; Oh, M. I.; Swanson, J. M. J., Improving the accuracy and convergence of drug permeation simulations via machine-learned collective variables. J. Chem. Phys. 2021, 155 (4), 045101.

48. Abraham, M. J.; Murtola, T.; Schulz, R.; Páll, S.; Smith, J. C.; Hess, B.; Lindahl, E., GROMACS: High performance molecular simulations through multi-level parallelism from laptops to supercomputers. SoftwareX 2015, 1-2, 19–25.

49. Tribello, G. A.; Bonomi, M.; Branduardi, D.; Camilloni, C.; Bussi, G., PLUMED 2: New feathers for an old bird. Comput. Phys. Commun. 2014, 185 (2), 604–613.

50. Klauda, J. B.; Venable, R. M.; Freites, J. A.; O’Connor, J. W.; Tobias, D. J.; Mondragon-Ramirez, C.; Vorobyov, I.; MacKerell, A. D.; Pastor, R. W., Update of the CHARMM All-Atom Additive Force Field for Lipids: Validation on Six Lipid Types. J. Phys. Chem. B 2010, 114 (23), 7830–7843.

51. Vanommeslaeghe, K.; Hatcher, E.; Acharya, C.; Kundu, S.; Zhong, S.; Shim, J.; Darian, E.; Guvench, O.; Lopes, P.; Vorobyov, I.; Mackerell Jr., A. D., CHARMM general force field: A force field for drug-like molecules compatible with the CHARMM all-atom additive biological force fields. J. Comput. Chem. 2010, 31 (4), 671–690.

52. Jorgensen, W. L.; Chandrasekhar, J.; Madura, J. D.; Impey, R. W.; Klein, M. L., Comparison of simple potential functions for simulating liquid water. J. Chem. Phys. 1983, 79 (2), 926–935.

53. Jo, S.; Lim, J. B.; Klauda, J. B.; Im, W., CHARMM-GUI Membrane Builder for Mixed Bilayers and Its Application to Yeast Membranes. Biophys. J. 2009, 97 (1), 50–58.

54. Martínez, L.; Andrade, R.; Birgin, E. G.; Martínez, J. M., PACKMOL: A package for building initial configurations for molecular dynamics simulations. J. Comput. Chem. 2009, 30 (13), 2157–2164.

55. Bussi, G.; Donadio, D.; Parrinello, M., Canonical sampling through velocity rescaling. J. Chem. Phys. 2007, 126 (1), 014101.

56. Berendsen, H. J. C.; Postma, J. P. M.; Gunsteren, W. F. v.; DiNola, A.; Haak, J. R., Molecular dynamics with coupling to an external bath. J. Chem. Phys. 1984, 81 (8), 3684–3690.

57. Essmann, U.; Perera, L.; Berkowitz, M. L.; Darden, T.; Lee, H.; Pedersen, L. G., A smooth particle mesh Ewald method. J. Chem. Phys. 1995, 103 (19), 8577–8593.

58. Hess, B.; Bekker, H.; Berendsen, H. J. C.; Fraaije, J. G. E. M., LINCS: A linear constraint solver for molecular simulations. J. Comput. Chem. 1997, 18 (12), 1463–1472.

59. Humphrey, W.; Dalke, A.; Schulten, K., VMD: Visual molecular dynamics. J. Mol. Graph. Model. 1996, 14 (1), 33–38.

60. Laio, A.; Parrinello, M., Escaping free-energy minima. Proc. Natl. Acad. Sci. U.S.A. 2002, 99 (20), 12562–12566.

61. Laio, A.; Gervasio, F. L., Metadynamics: a method to simulate rare events and reconstruct the free energy in biophysics, chemistry and material science. Rep. Prog. Phys. 2008, 71 (12), 126601.

62. E W.; Ren, W.; Vanden-Eijnden E., Finite Temperature String Method for the Study of Rare Events. J. Phys. Chem. B 2005, 109 (14), 6688–6693.

63. Cameron, M.; Kohn, R. V.; Vanden-Eijnden, E., The String Method as a Dynamical System. J. Nonlinear Sci. 2011, 21 (2), 193–230.

64. Sun, R.; Dama, J. F.; Tan, J. S.; Rose, J. P.; Voth, G. A., Transition-Tempered Metadynamics is a Promising Tool for Studying the Permeation of Drug-like Molecules through Membranes. J. Chem. Theory Comput. 2016, 12, 5157–5169.

65. Sun, R.; Han, Y.; Swanson, J. M. J.; Tan, J. S.; Rose, J. P.; Voth, G. A., Molecular transport through membranes: Accurate permeability coefficients from multidimensional potentials of mean force and local diffusion constants. J. Chem. Phys. 2018, 149 (7), 072310.

66. Naritomi, Y.; Fuchigami, S., Slow dynamics in protein fluctuations revealed by time-structure based independent component analysis: The case of domain motions. J. Chem. Phys. 2011, 134 (6), 065101.

67. Yang, Y. I.; Parrinello, M., Refining Collective Coordinates and Improving Free Energy Representation in Variational Enhanced Sampling. J. Chem. Theory Comput. 2018, 14 (6), 2889–2894.

68. Michaud-Agrawal, N.; Denning, E. J.; Woolf, T. B.; Beckstein, O., MDAnalysis: A toolkit for the analysis of molecular dynamics simulations. J. Comput. Chem. 2011, 32 (10), 2319–2327.

69. Barducci, A.; Bonomi, M.; Parrinello, M., Metadynamics. Wiley Interdiscip. Rev. Comput. Mol. Sci. 2011, 1 (5), 826–843.

70. E W.; Ren W.; Vanden-Eijnden, E., Simplified and improved string method for computing the minimum energy paths in barrier-crossing events. J. Chem. Phys. 2007, 126 (16), 164103.

71. Masoud, M. S.; Awad, M. K.; Shaker, M. A.; El-Tahawy, M. M. T., The role of structural chemistry in the inhibitive performance of some aminopyrimidines on the corrosion of steel. Corros. Sci. 2010, 52 (7), 2387–2396.

72. Exner, O.; Jehlička, V., Conformation around several equivalent bonds. Polymethoxy derivatives of benzene. Collect. Czech. Chem. Commun. 1983, 48 (4), 1030–1041.

73. Mehdi, S.; Wang, D.; Pant, S.; Tiwary, P., Accelerating All-Atom Simulations and Gaining Mechanistic Understanding of Biophysical Systems through State Predictive Information Bottleneck. J. Chem. Theory Comput. 2022, 18 (5), 3231–3238.

74. Bonati, L.; Piccini, G.; Parrinello, M., Deep learning the slow modes for rare events sampling. Proc. Natl. Acad. Sci. U.S.A. 2021, 118 (44), e2113533118.

75. Chen, H.; Chipot, C., Chasing collective variables using temporal data-driven strategies. QRB Discov. 2023, 4, e2.

76. Shmilovich, K.; Ferguson, A. L., Girsanov Reweighting Enhanced Sampling Technique (GREST): On-the-Fly Data-Driven Discovery of and Enhanced Sampling in Slow Collective Variables. J. Phys. Chem. A 2023, 127 (15), 3497–3517.

77. Donati, L.; Keller, B. G., Girsanov reweighting for metadynamics simulations. J. Chem. Phys. 2018, 149 (7).

