## Supporting Information for "tICA-Metadynamics for Identifying Slow Dynamics in Membrane Permeation"

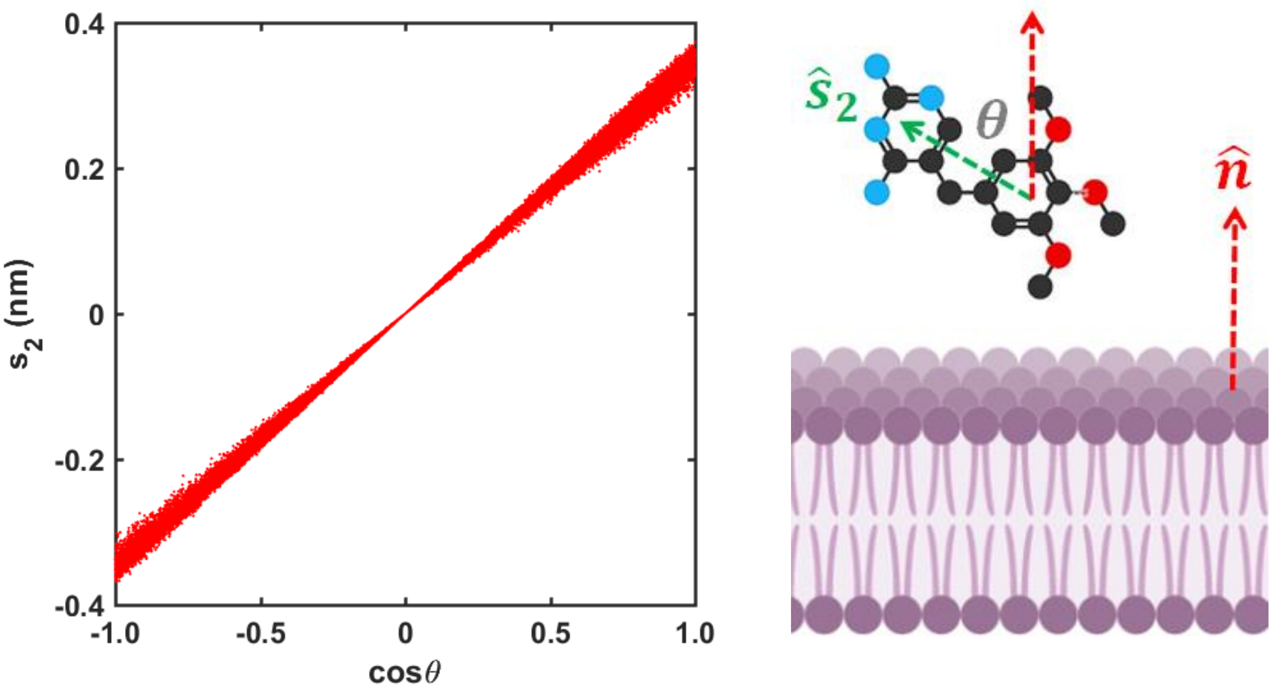


**Figure S1.** Orientational CV $s_2$ as a function of $cos\,\theta$ where $\theta$ is the angle between the unit vector $\hat{s}_2$ (green dotted arrow) of trimethoprim and the surface normal $\hat{n}$ (red dotted arrow) of the nearest membrane leaflet.

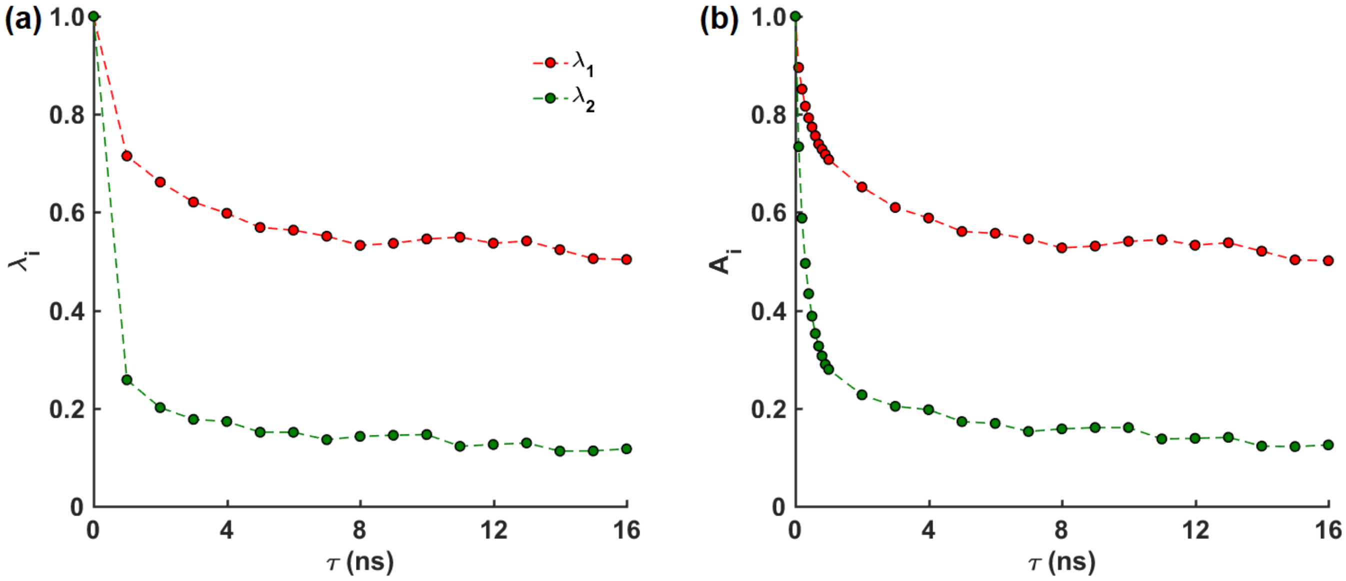


**Figure S2.** (a) Eigenvalues $\lambda_i$ corresponding to the first (red, $i = 1$) and second (green, $i = 2$) eigenvectors obtained from tICA on the reweighted trajectory. (b) Autocorrelations $A_i$ of the projections of the data points onto the first (red, $i = 1$) and second (green, $i = 2$) eigenvectors obtained from tICA.

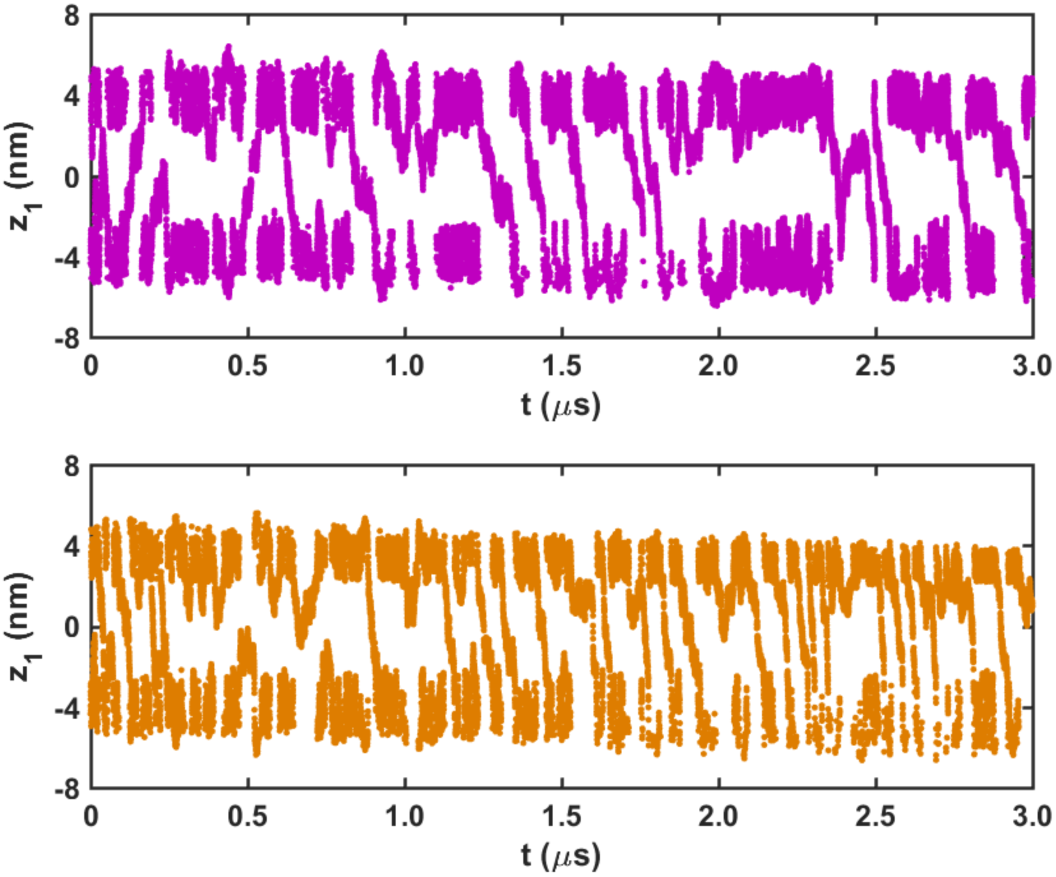


**Figure S3.** Time evolution of $z_1$ when the suboptimal CV $s_0$ (purple, upper panel) and the optimal CVs $s_1$ and $s_2$ (orange, lower panel) are biased.

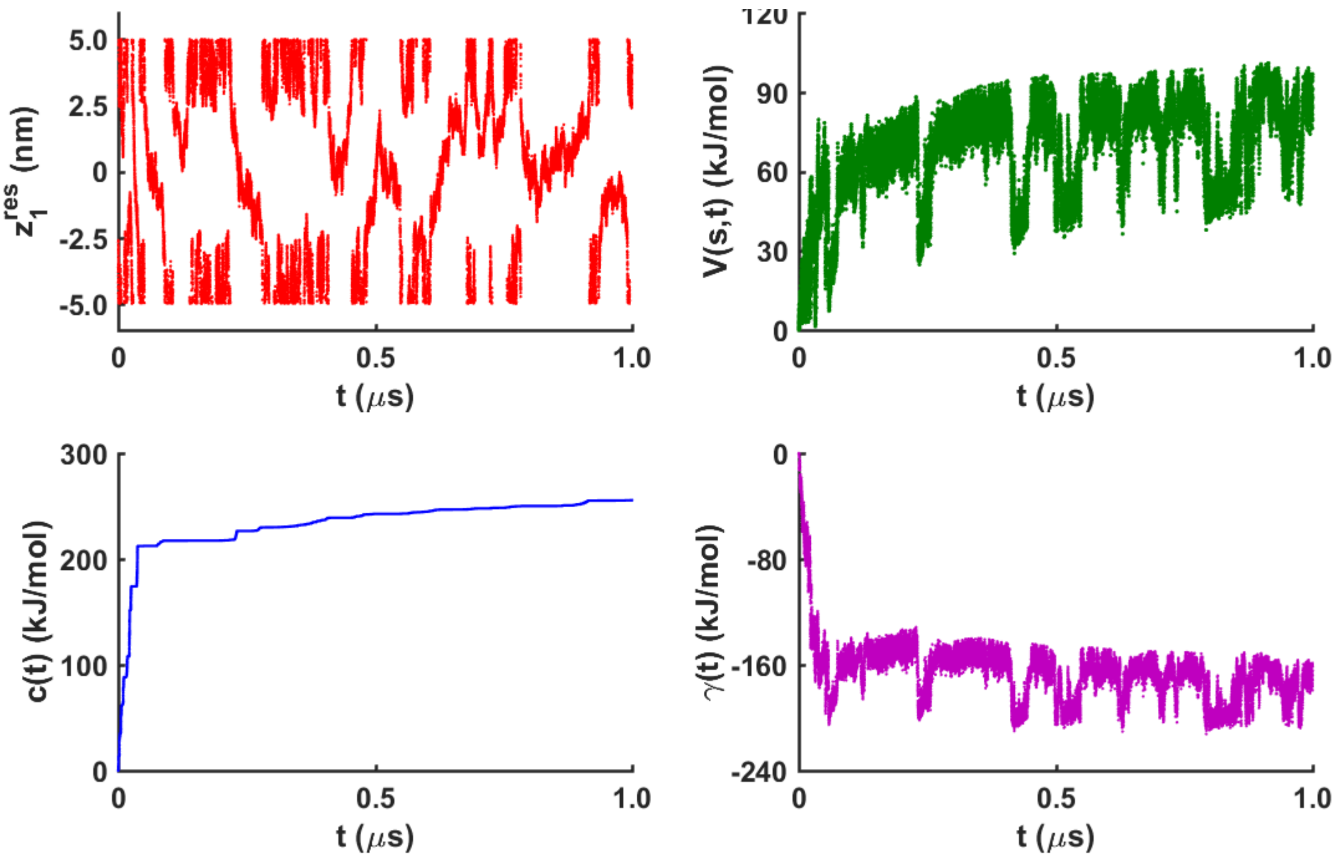


**Figure S4.** Time evolution of rescaled $z_1$ (red, upper left panel), (2) the instantaneous potential $V(s,t)$ (green, upper right panel), (3) the bias offset $c(t)$ (blue, lower left panel), and (d) the difference $\gamma(t)$ (purple, lower right panel) between $V(s,t)$ and $c(t)$ in the 1 $\mu s$-long TTMetaD simulation we performed.

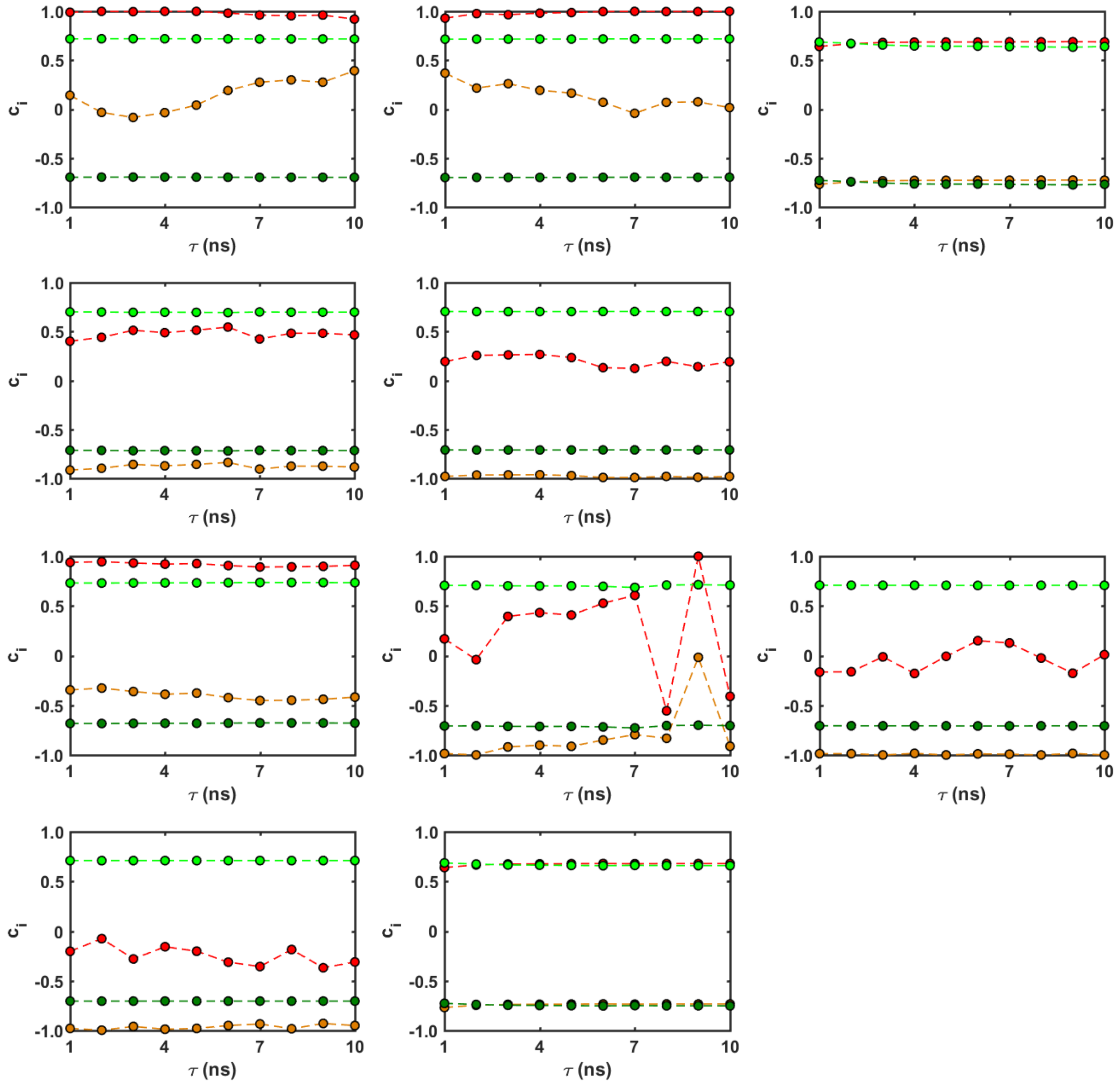


**Figure S5.** tICA results from 10 replicas of the WTMetaD trajectories with the PBCs. The same color scheme was used as in Fig. 4(a-b) to represent the components of the first and second eigenvectors.

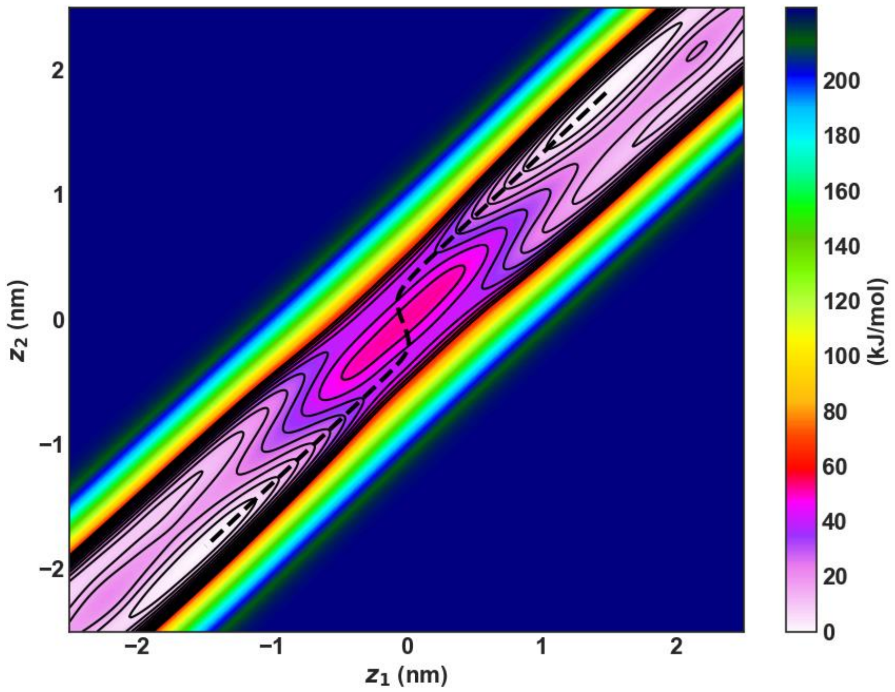


**Figure S6.** Average 2D PMF along the tICA CVs $z_1$ and $z_2$ in the presence of the harmonic walls. The dashed black line indicates the MFEP found from the zero-temperature string method.

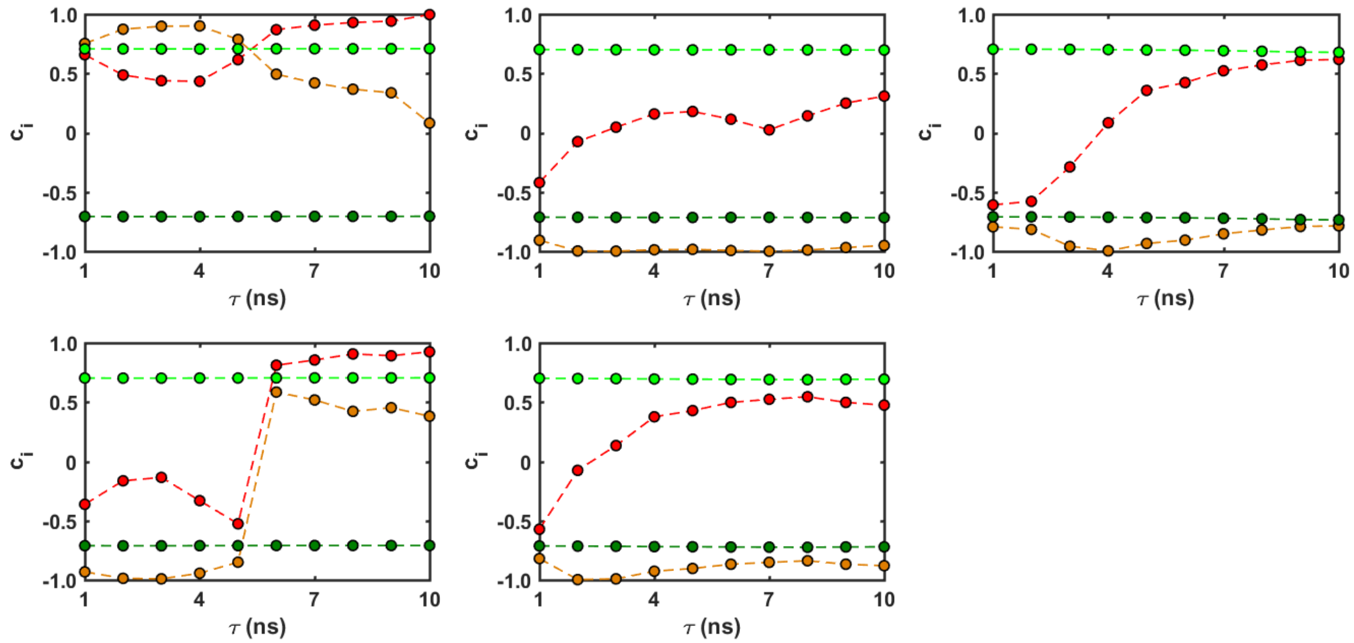


**Figure S7.** tICA results from 5 replicas of the WTMetaD trajectories with the harmonic walls. The same color scheme was used as in Fig. 4(a-b) to represent the components of the first and second eigenvectors.

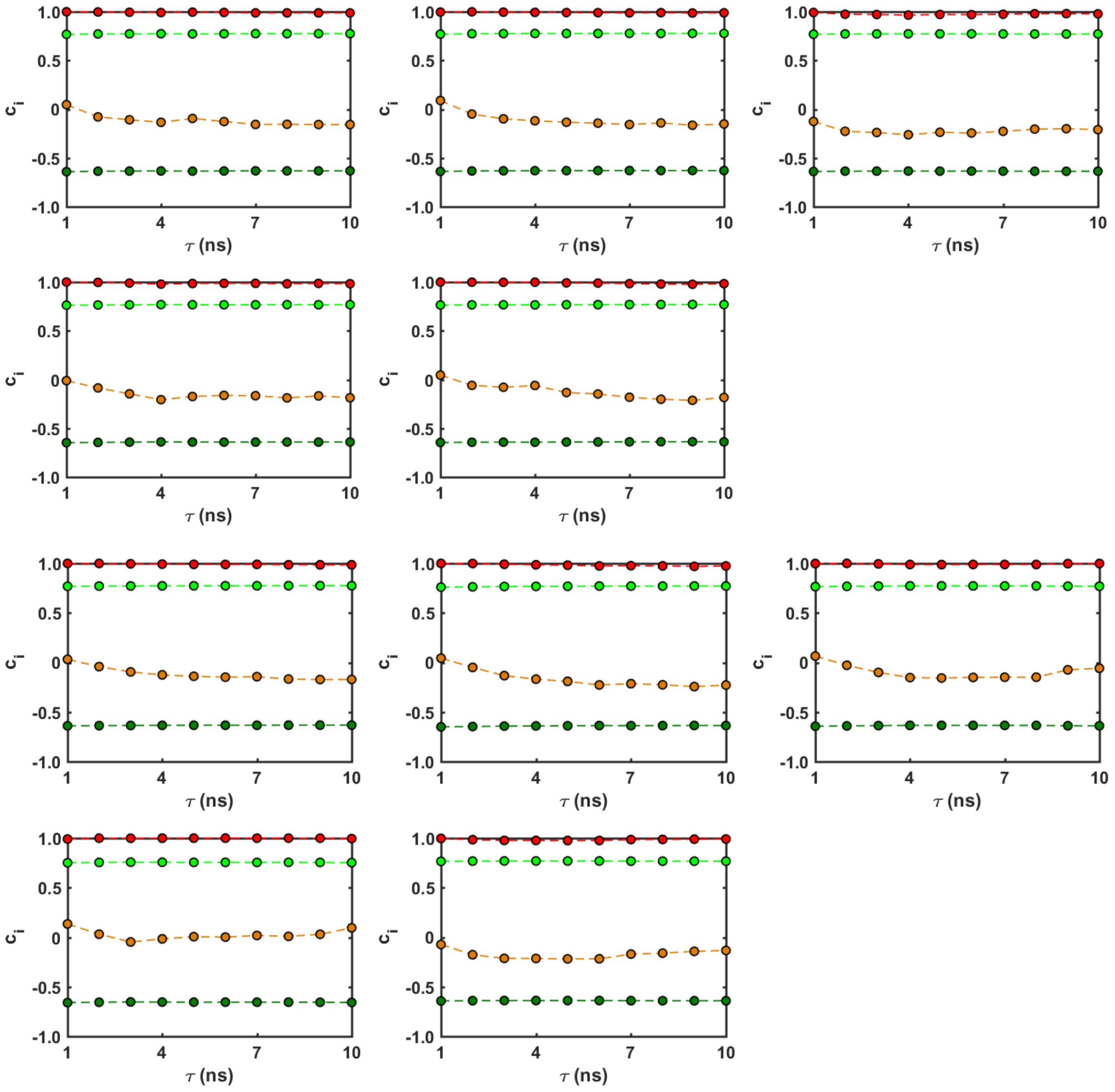


**Figure S8.** tICA results from 10 replicas of the WTMetaD trajectories with the absolute values of the molecular features. The same color scheme was used as in Fig. 4(a-b) to represent the components of the first and second eigenvectors.

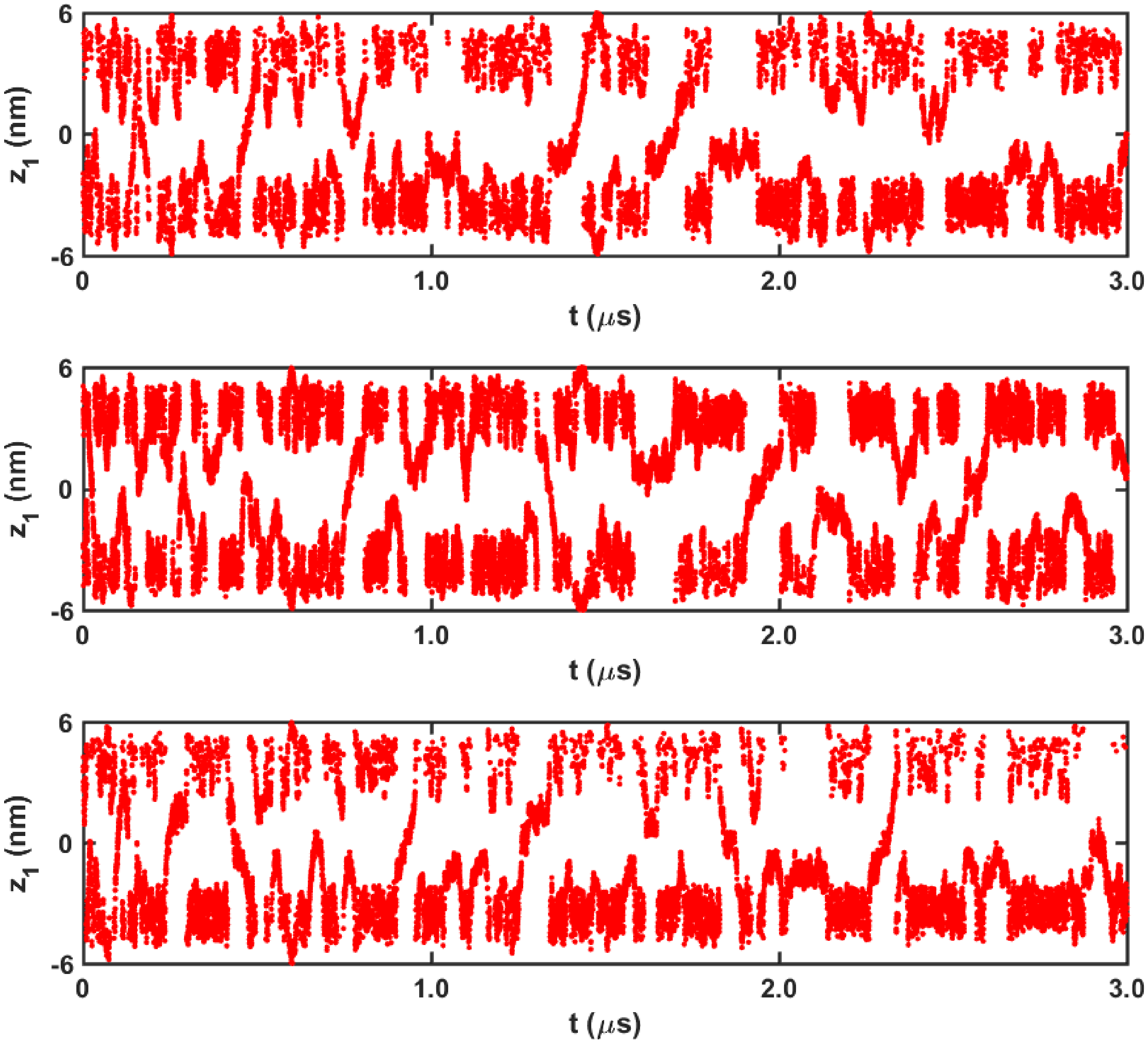


**Figure S9.** Time evolution of $z_1$ in three WTMetaD simulations where $s_1'$ and $s_2'$ are biased.

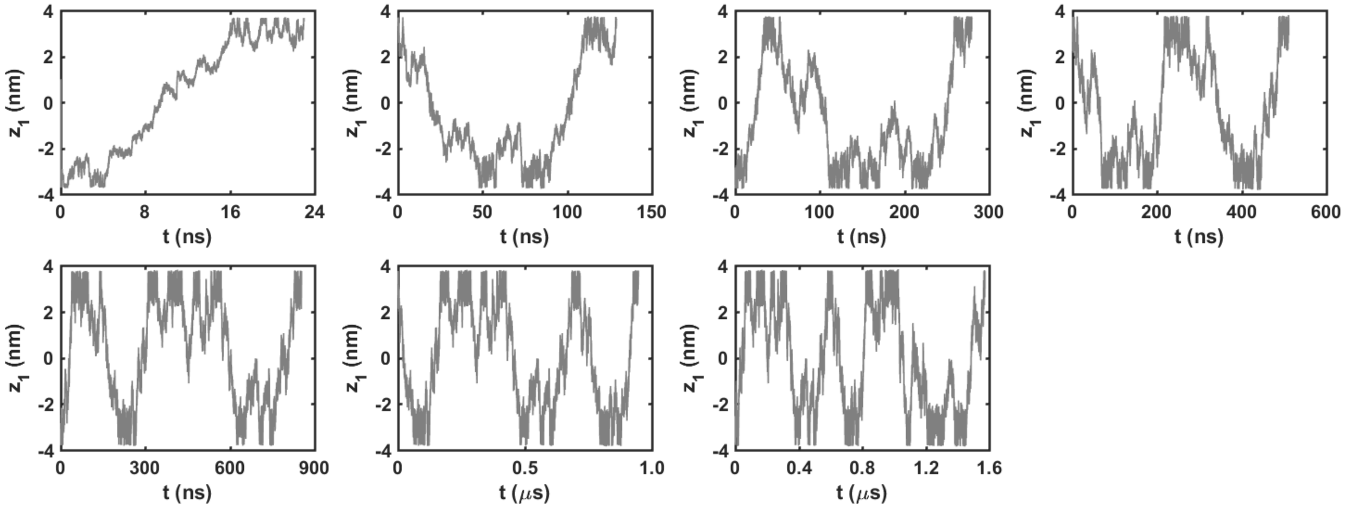


**Figure S10.** Examples of subtrajectories involving one to seven rare events (*i.e.*, membrane crossings of trimethoprim).

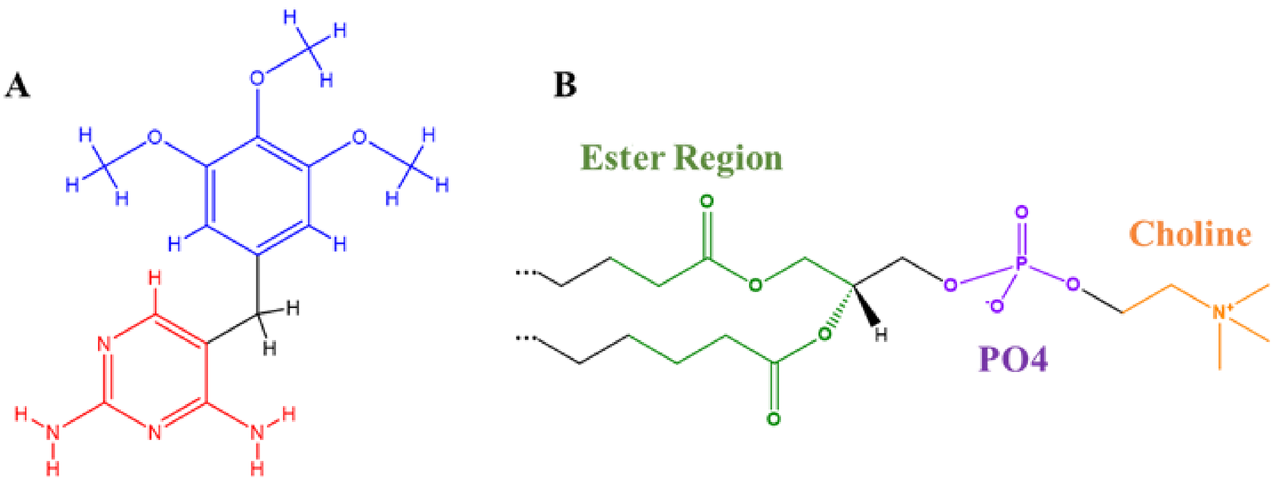


**Figure S11.** (A) DAP (red) and TMB (blue) groups in trimethoprim; (B) Ester (green), phosphate (purple), and choline (orange) groups of a POPC molecule for the contact and interaction energy analyses. Full tails are not shown for better visualization.

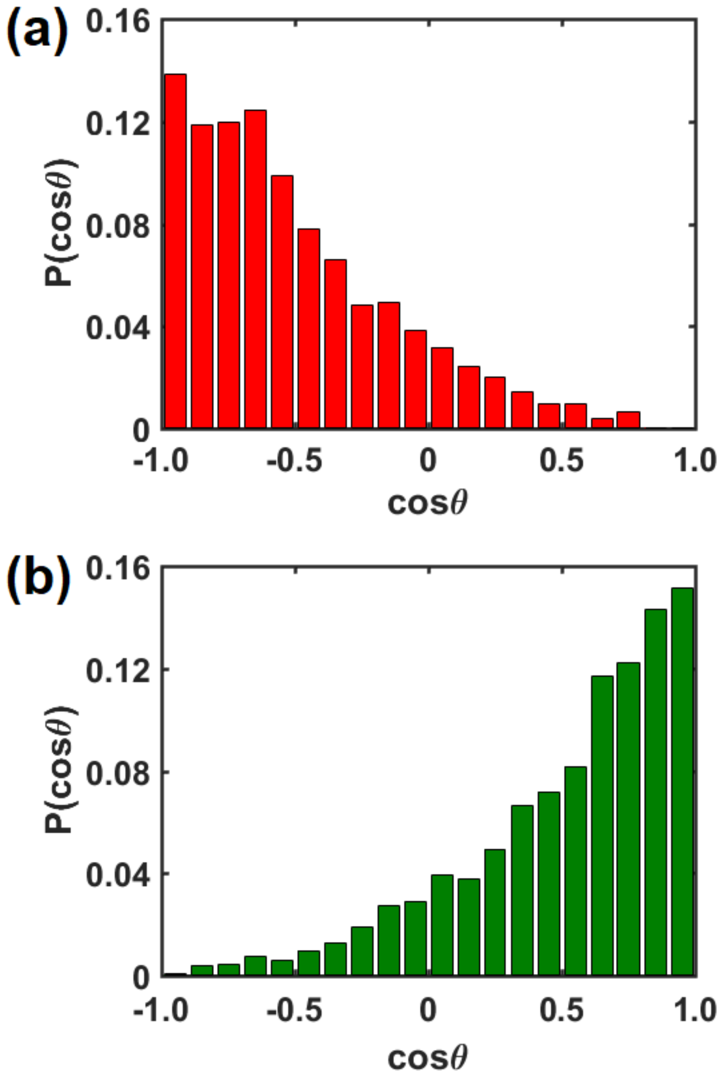


**Figure S12**. Probability distributions of the orientation angle $\theta$ of trimethoprim with respect to the surface normal of the nearest leaflet (a) when it is very close to and above the membrane surface (in the aqueous region) and (b) when it resides inside the membrane (at the metastable states) in the 5 $\mu$s-long unbiased MD simulation.

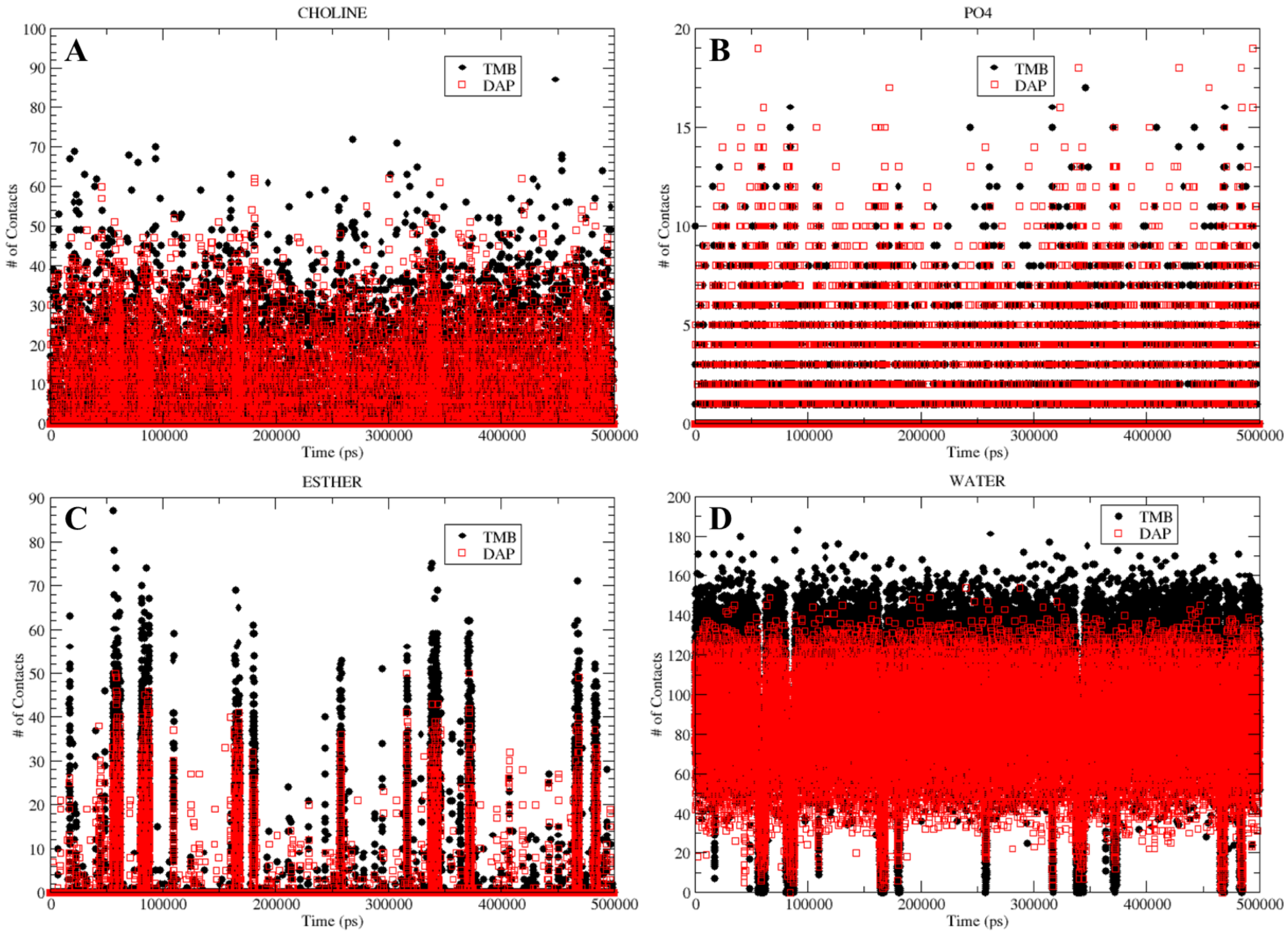


**Figure S13.** Number of contacts of TMB (black filled sphere) and DAP (red square) with POPC components: (A) choline; (B) phosphate; (c) ester region; and with (D) water molecules in the 5 $\mu$s-long unbiased simulation.

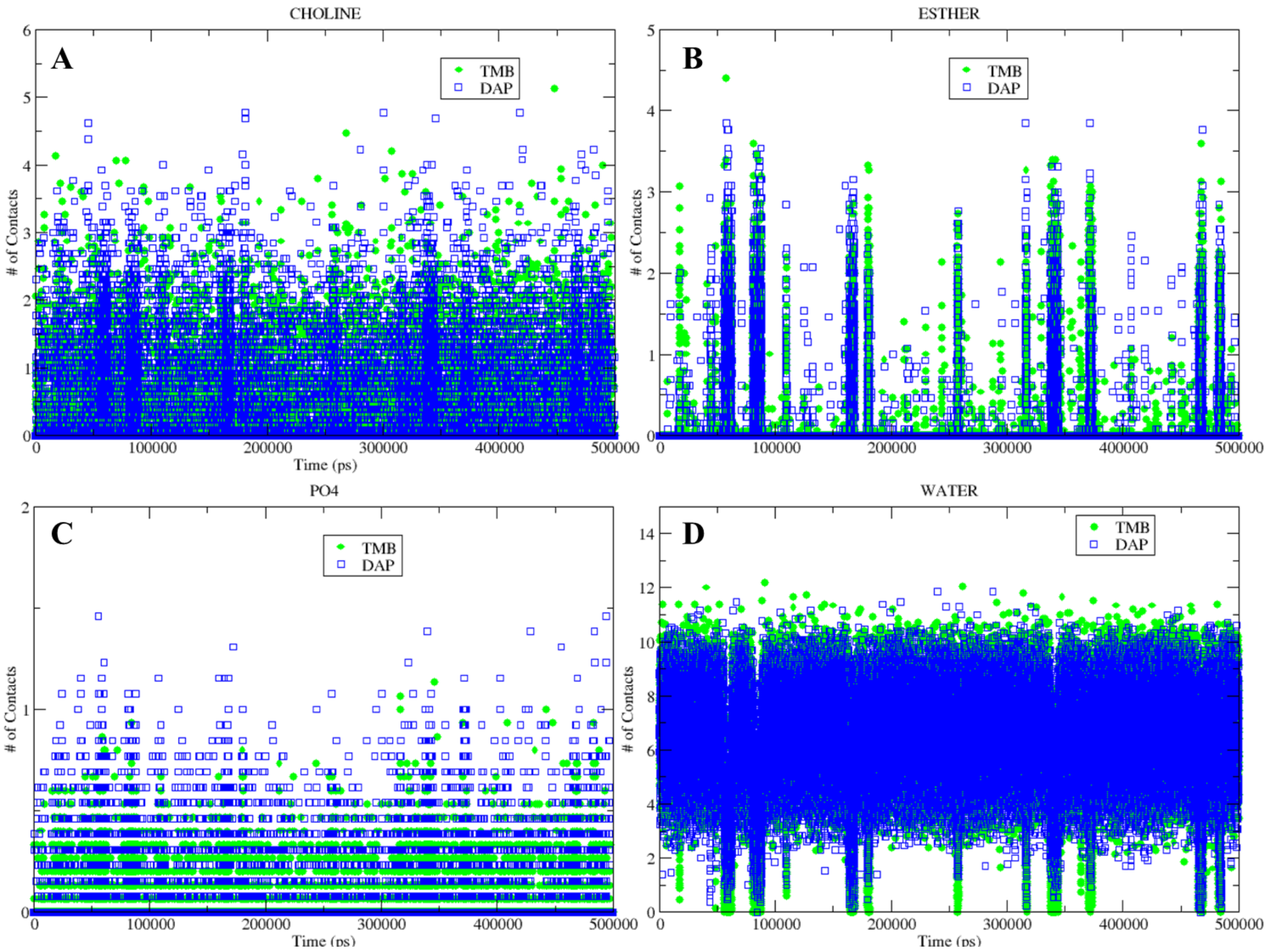


**Figure S14.** Number of contacts per atom of TMB (green filled sphere) and DAP (blue square) with POPC components: (A) choline; (B) ester region; (C) phosphate; and with (D) water molecules in the 5 $\mu$s-long unbiased simulation. For consistency, all atoms of the DAP group (13) and the atoms of the TMB methoxy groups and its C from the ring (15) were selected for this analysis.

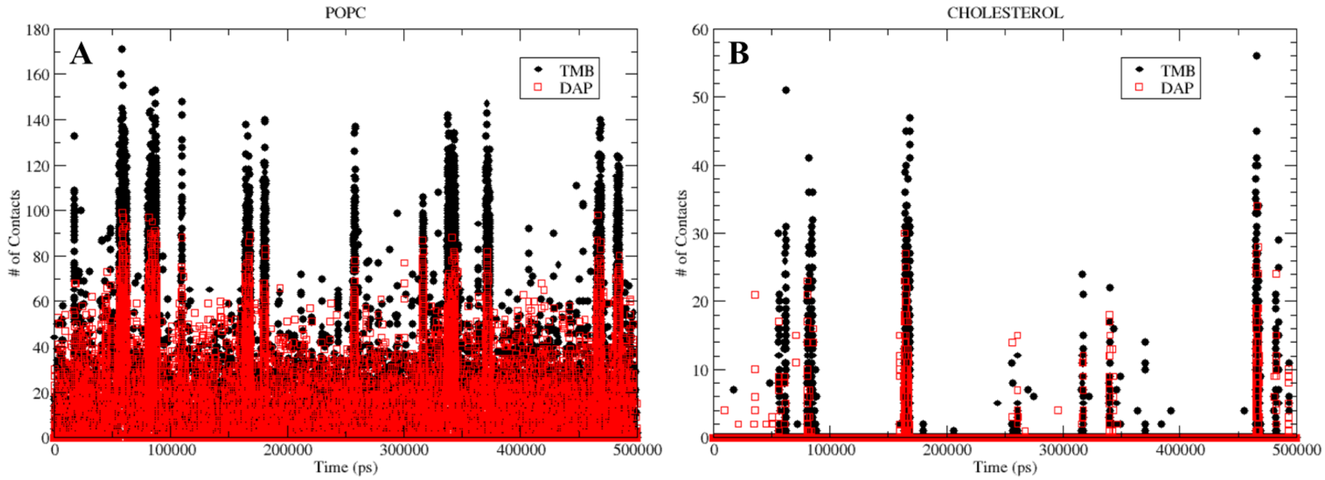


**Figure S15.** Number of contacts of TMB (black filled sphere) and DAP (red square) with (A) POPC and (B) cholesterol in the 5 $\mu$s-long unbiased simulation.

**Table S1.** Average interaction energy with standard error (in kJ/mol) and relative increase (RI, %) for TMB/DAP groups interacting with each component of the system and POPC regions.

| Groups | Interaction energy (kJ/mol) | | | | | |
|---|---|---|---|---|---|---|
| | Membrane | POPC | CHL | $PO_4$/Choline | Ester | Water |
| DAP | -10.28 ± 0.12 | -10.17 ± 0.12 | -0.11 ± 0.01 | -7.08 ± 0.08 | -2.21 ± 0.04 | -134.00 ± 0.13 |
| TMB | -13.69 ± 0.20 | -13.45 ± 0.20 | -0.24 ± 0.01 | -7.74 ± 0.17 | -4.34 ± 0.07 | -139.93 ± 0.25 |
| RI§ | 24.9 | 24.4 | 54.2 | 8.5 | 49.1 | 4.2 |

§RI = [(TMB – DAP)/TMB * 100]

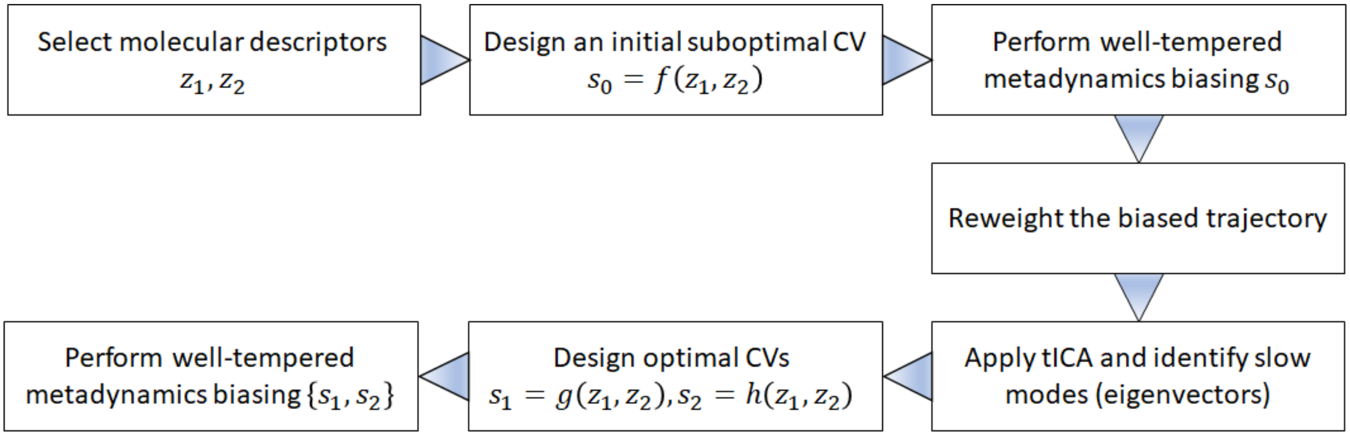


**Figure S116.** Schematic diagram that summarizes the key steps involved in the tICA-MetaD protocol.
